# Loss of cilia after neurulation impacts brain development and neuronal activity in larval zebrafish

**DOI:** 10.1101/2023.09.20.558654

**Authors:** Percival P. D’Gama, Inyoung Jeong, Andreas Moe Nygård, Anh-Tuan Trinh, Emre Yaksi, Nathalie Jurisch-Yaksi

## Abstract

Cilia are slender, hair-like structures extending from cell surfaces and playing essential roles in diverse physiological processes. Within the nervous system, primary cilia contribute to signaling and sensory perception, while motile cilia facilitate cerebrospinal fluid flow. Here, we investigated the impact of ciliary loss on neural circuit development using a zebrafish line displaying ciliogenesis defects. We found that cilia loss after neurulation affects neurogenesis and brain morphology, and lead to altered gene expression profiles. Using whole brain calcium imaging, we measured reduced light-evoked and spontaneous neuronal activity in all brain regions. By shedding light on the intricate role of cilia in neural circuit formation and function in the zebrafish, our work highlights their evolutionary conserved role in the brain and set the stage for future analysis of ciliopathy models.

## INTRODUCTION

Cilia are hair-like structures that extend from the surface of cells (Mitchison and Valente, 2017; Nachury, 2014). They play a vital role in various physiological processes across species (Louvi and Grove, 2011). In the context of the nervous system, cilia are involved in functions ranging from cellular signaling to sensory perception and movement of cerebrospinal fluid (CSF) (Bachmann-Gagescu and Neuhauss, 2019; Bear and Caspary; Falk et al., 2015; Guemez-Gamboa et al., 2014; Ringers et al., 2020; Suciu and Caspary, 2021).

Primary cilia are present on neuronal progenitors and have been described in many species, including human, mouse, and zebrafish (Gabriel et al., 2016; Hansen et al., 2021; Paridaen and Huttner, 2014). They serve as essential signaling hubs which receive and transmit cues (Wachten and Mick, 2021), such as hedgehog signaling (Bangs and Anderson, 2017; Huangfu et al., 2003). Cilia intricately regulate proliferation and differentiation (Gabriel et al., 2016; Youn and Han, 2018), patterning (Higginbotham et al., 2013), migration (Higginbotham et al., 2012; Stoufflet and Caillé, 2022; Stoufflet et al., 2020), axon guidance (Guo et al., 2019; Suciu and Caspary, 2021), synaptogenesis (Kumamoto et al., 2012) and connectivity (Guo et al., 2015; Guo et al., 2017). Giving their ubiquitous function, cilia-related developmental defects have been described in multiple brain structures including the cortex, hippocampus, and cerebellum in mammals (Liu et al., 2021; Suciu and Caspary, 2021).

In differentiated neurons, primary cilia play a dual role. They can participate in sensing sensory modalities such as odors (Bergboer et al., 2018; Falk et al., 2015; McClintock et al., 2020) and light (Bachmann-Gagescu and Neuhauss, 2019; Bujakowska et al., 2017; Insinna and Besharse, 2008), or they can actively influence neuronal physiology (DeMars et al., 2023; Sheu et al., 2022; Wang et al., 2021). For example, cilia were shown to regulate food uptake through the modulation of hypothalamic neurons in the paraventricular nucleus (Wang et al., 2021) and circadian rhythm through interneuronal coupling in the super chiasmatic nucleus (Tu et al., 2023). Cilia can also alter the chromatin accessibility of hippocampal neurons through axo-ciliary serotonergic synapses (Sheu et al., 2022). Ciliary signaling usually involves G protein-coupled receptors, which localize to the primary cilium, such as olfactory receptors, somatostatin receptor type 3 (SSTR3), serotonin receptor 6 (HTR6), melanin-concentrating hormone receptor 1 (MCHR1), or dopamine receptor 1 (D1) (Berbari et al., 2008; Domire et al., 2011; Hamon et al., 1999; Hilgendorf et al., 2016; Händel et al., 1999; Menco et al., 1997; Sengupta et al., 1996; Sheu et al., 2022; Wachten and Mick, 2021).

In contrast to primary cilia, motile cilia propel fluids and particles across tissue surfaces (Fliegauf et al., 2007; Reiten et al., 2017). In the brain, cilia-driven fluid flow is important for the movement and homeostasis of CSF (D’Gama et al., 2021; Del Bigio, 2010; Faubel et al., 2016; Olstad et al., 2019; Ringers et al., 2020; Sawamoto et al., 2006). Intriguingly, CSF flow has also been implicated in neuronal migration in mice, through the potential establishment of morphogen gradients (Sawamoto et al., 2006).

Human patients with dysfunctional cilia can present several neurodevelopmental and neurological symptoms (Andreu-Cervera et al., 2021; Reiter and Leroux, 2017; Suciu and Caspary, 2021). For instance, mutations in the genes encoding Bardet-Biedl syndrome (BBS) proteins are associated with retinal degeneration, hyperphagia and learning disabilities (Forsyth and Gunay-Aygun, 1993; Forsythe and Beales, 2013). Another classical example is Joubert syndrome (Bachmann-Gagescu et al., 2020; Joubert et al., 1969; Parisi and Glass, 1993), which is a rare genetic disorder affecting the development of the cerebellum and brainstem (Brancati et al., 2010; Ferland et al., 2004). Joubert syndrome is characterized by cerebellar vermis malformation, disrupting motor coordination in addition to cognitive dysfunction and epilepsy (Bachmann-Gagescu et al., 2020; Bachmann-Gagescu et al., 2015). Altogether, these findings suggest that disruptions in ciliary function impact neuronal development, which may influence neuronal activity in the brain and even lead to epileptic seizures. However, it is still unclear how primary cilia loss affects the establishment of neuronal circuits in the brain and whether it leads to altered neural activity.

In this study, we leveraged the small size, transparency, and genetic amenability of zebrafish larvae to investigate the impacts of ciliary loss on the development of neural circuits. To this end, we used a cilia mutant carrying a mutation in the ciliary gene *traf3ip1* (also known as *ift54* or *elipsa*), which was shown to abolish all cilia at larval stages (Omori et al., 2008). In our study, we first determined the onset of ciliary defects in the *elipsa* mutant. We identified a progressive ciliary loss between the 10 somites stage and 30 hours post-fertilization (hpf), allowing us to study specifically the role of cilia after the process of neurulation. Upon histological analysis, we observed that ciliary loss alters the overall brain morphology of zebrafish larvae and increases cell proliferation in the optic tectum, which is the visual processing center in zebrafish. Next, using transcriptomics, we revealed that genes and pathways involved in retinal function and neuronal development, including hedgehog signaling, are dampened by the loss of cilia. Finally, we identified that the mutants have reduced light-evoked and ongoing neuronal activity in all brain areas, but do not display seizures. Taken together, our work identified that cilia are critical for brain development and physiology of neural circuits in zebrafish. This sets the stage for future analysis of ciliopathy models.

## RESULTS

### Loss-of-function mutation of *traf3ip1* (*elipsa*) abolishes cilia in the developing brain following neurulation

To study the impact of ciliary loss on brain development, we selected a zebrafish mutant line, *elipsa* or *traf3ip1tp49d* (Omori et al., 2008), which was shown to lack all primary and motile cilia at larval stages. We first aimed to identify the onset and penetrance of cilia loss in the *elipsa* mutant brain upon immunostaining of various developmental stages from 10 somites to 4 days post-fertilization (dpf) larvae. We used antibodies against acetylated tubulin and the ciliary protein arl13b, which labels both primary and motile cilia (Duldulao et al., 2009). To label motile cilia specifically, we used glutamylated tubulin as a marker as previously described (D’Gama and Jurisch-Yaksi, 2023; D’Gama et al., 2021; Olstad et al., 2019). Upon staining with anti-acetylated tubulin at 10 somites stage, we found no major differences in cilia between control and mutants in the neural keel, developing optic vesicles **(figure S1A1, A2 1B1, B2)** and in the left-right organizer (Kupffer’s vesicle) **(figure S1A3 1B3)**, suggesting that the maternal contribution of the gene prevents a ciliary phenotype at these early developmental stages.

Next, we investigated later time points of development following neurulation, from 30 hpf up to 4 dpf, when many mutants were still healthy and did not show major abnormalities beside their curved body axis and a mild heart oedema. We noticed a total loss of cilia in the *elipsa* embryos at 30 hpf **(figure S2A1-B4)**, which was maintained at 2 dpf **(figure S2C1-F4)** and 4 dpf **(figure 1)**. Notably, we observed that both primary cilia **(figure 1A1-B4)** and motile cilia **(figure 1C1-D4)** were absent in the entire brain at 2 dpf and 4 dpf. Taken together, our results reveal a progressive loss of cilia in the brain of *elipsa* mutant after neurulation, allowing us to study the impacts of cilia on the development and physiology of the nervous system in an animal devoid of early neural tube defects.

**Figure 1:**
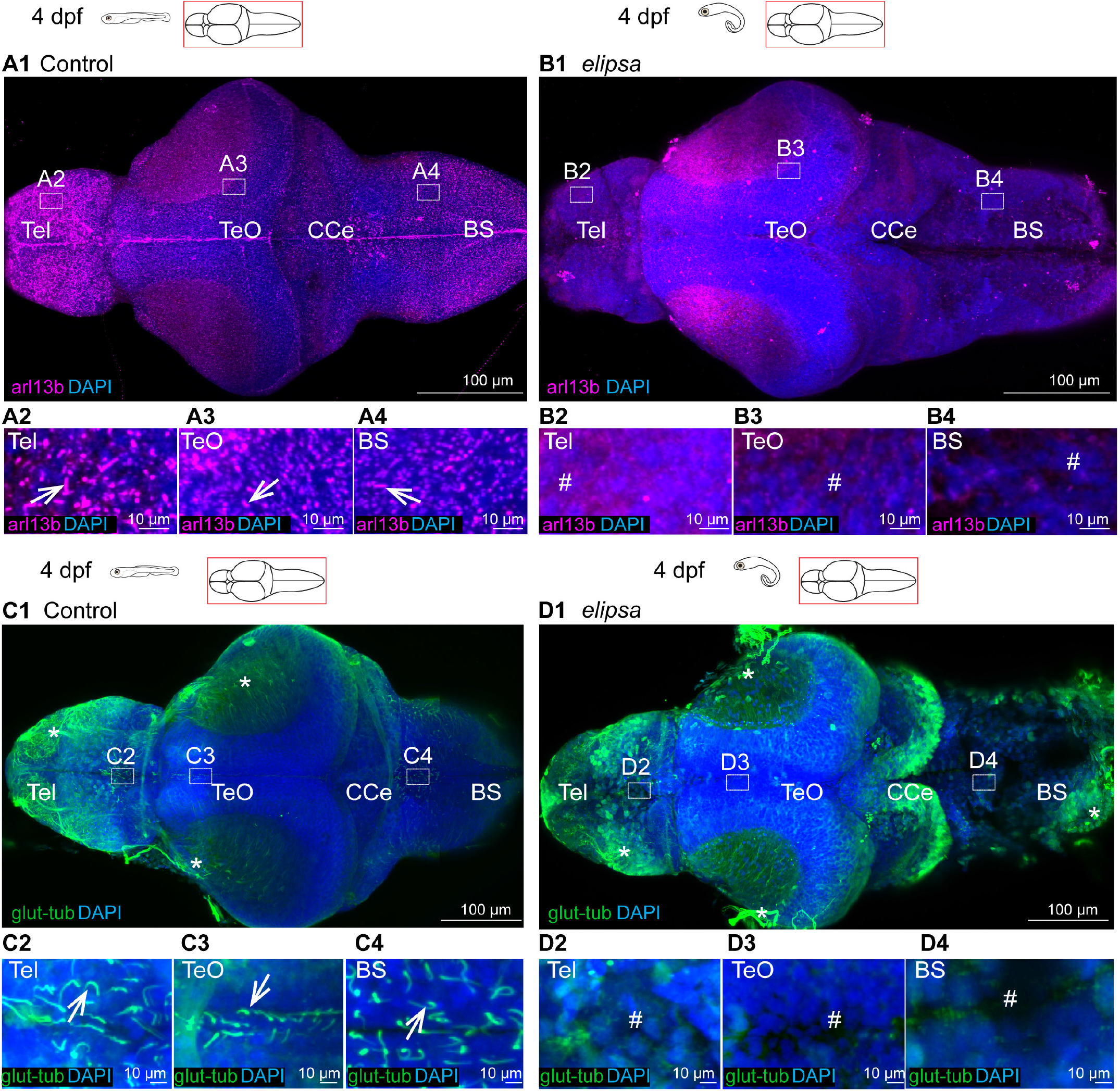
Loss of primary and motile cilia in the *elipsa* mutant larval brain. **(A1-A4** and **B1-B4)** Staining of dissected 4 dpf brains with arl13b antibody to stain all cilia in the brain, n=3. A1 At 4 dpf, arl13b stained cilia were located all over the brain, represented in insets drawn in the telencephalon **(A2)**, optic tectum **(A3)** and brainstem **(A4). (B1)** In the *elipsa* mutants, cilia were absent in the entire brain, as represented by insets drawn in the telencephalon **(B2)**, optic tectum **(B3)**, and brain stem **(B4). (C1-C4** and **D1-D4)** Staining of dissected 4 dpf brains with glutamylated tubulin used to identify motile cilia, n=5. **(C1)** At 4 dpf, single glutamylated tubulin-positive cilia were present in the forebrain choroid plexus **(C2)**, on the dorsal roof and ventral part **(C3)** of the tectal/diencephalic ventricle and in the rhombencephalic choroid plexus **(C4). (D1)** In *elipsa* mutant, glutamylated tubulin-positive cilia were absent in the telencephalon **(D2)**, optic tectum **(D3)**, and brain stem **(D4)**. Tel, Telencephalon; Teo, Optic Tectum; CCe, Corpus Cerebelli; BS, Brain stem. Cilia loss indicated by # symbol and nonspecific signal from glutamylated tubulin is represented by * symbol.

### Cilia loss after neurulation leads to abnormal brain morphology and increases proliferation in the optic tectum

To identify the impact of cilia loss on brain development, we conducted various estimations of the brain size, including the width, length, and height of the telencephalon **(figure 2A1-A4)**, optic tectum **(figure 2B1-B4)**, and hindbrain **(figure 2C1-C4)**. In our analysis we found a significant decrease in the length of the telencephalon **(figure 2A3)**, the width of the optic tectum **(figure 2B2)**, cerebellum **(figure 2C2)** and brain stem **(figure 2C3)**. In contrast, the *elipsa* mutants displayed a significant increase in hindbrain length **(figure 2C4)**. Interestingly, while the overall width of the optic tectum was reduced, the thickness and density of the cellular layer in the tectum was increased **(figure 2D2, D4)**. Notably, malformation of the hindbrain, especially the cerebellum, was highly apparent upon staining with glutamylated tubulin at 4 dpf **(figure 1C1-D1)**.

**Figure 2:**
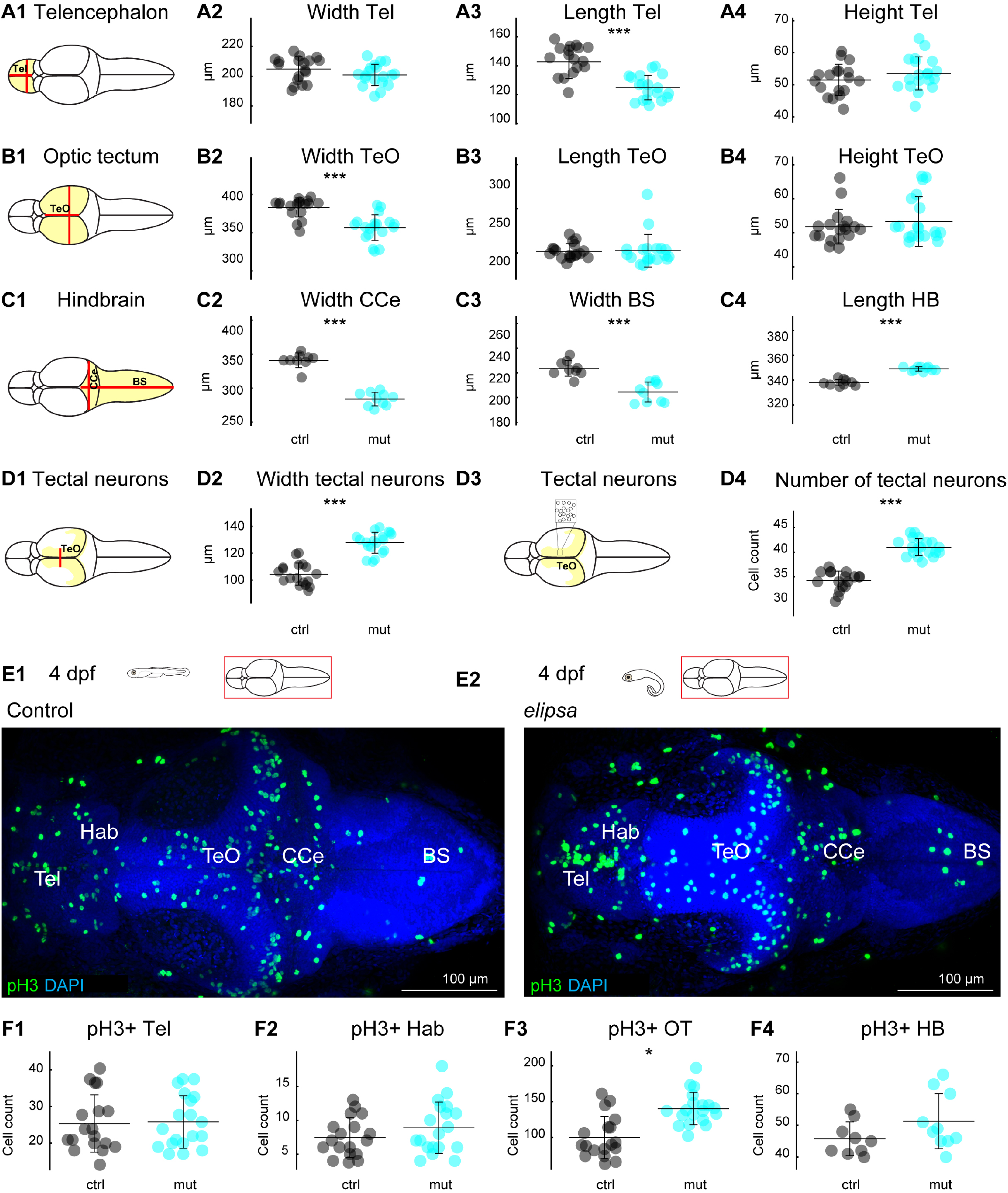
Cilia loss leads to abnormal brain size and altered cell proliferation in the optic tectum. **(A1-D4)** Quantification of brain morphology of 4 dpf larval brains for control (black) and *elipsa* mutants (cyan). Schematic representation of the measurements for **(A1)** telencephalon, **(B1)** optic tectum, **(C1)** hindbrain regions and **(D1)** tectal neurons. Brain size was estimated by measuring the width, length, and height of telencephalon **(A2, A3, A4)**, optic tectum **(B2, B3, B4)**, width of CCe **(C2)**, width of BS **(C3)**, length of hindbrain **(C4)**, and the width **(D2)** and number **(D4)** of tectal neurons. Controls are in black and *elipsa* mutants in cyan. n =18 controls and 19 mutants for **A2, A3, A4, B2, B3, B4, D2, D4**. n = 9 controls and 10 mutants for **C2, C3** and **C4. (E1, E2)** staining for mitotic cells using an anti-pH3 antibody. **(F1-F4)** Cell count for pH3 positive cells (pH3+) in telencephalon **(F1)**, habenula **(F2)**, optic tectum **(F3)** and hindbrain **(F4)**. *: p < 0.05, ***: p < 0.001 according to Wilcoxon Rank Sum test. Mean+/- SD (standard deviation) is indicated on scatter plots. Tel, Telencephalon; Teo, Optic Tectum; BS, Brain stem; CCe, Corpus Cerebelli; HB, Hindbrain.

To determine if the defects in brain size were due to a reduction in cell proliferation, we stained mitotic cells using a phosphorylated Histone H3 (pH3) antibody (**figure 2E1-E2**) (Prigent and Dimitrov, 2003). We then quantified pH3 positive cells in distinct brain regions **(figure 2F1-F4)**. Our quantifications showed a significantly higher number of mitotic cells in the optic tectum in *elipsa* mutants **(figure 2F3)** while we did not observe differences in cell numbers in other brain regions **(figure 2F1, F2 and F4)**. These results are consistent with our observations of increased cell density in the optic tectum in the *elipsa* mutants **(figure 2D4)**. Our findings suggest that loss of cilia after neurulation disrupts normal brain development and results in an overall smaller brain with more pronounced malformations in the optic tectum and hindbrain.

### Transcriptomic analysis identifies differentially regulated genes involved in phototransduction and brain development

To explore the impact of cilia loss on gene expression, we performed RNA-sequencing of whole 4 dpf larvae. We identified a total of 520 differentially expressed genes (DEGs), comprising 100 upregulated DEGs and 420 downregulated DEGs **(figure 3A1-A2)**, between control and *elipsa* larvae. We next classified the genes into different Gene Ontology (GO) terms consisting of GO biological process, GO cellular component, and GO molecular function **(figure 3B1-B3)**. Remarkably, the prominent GO biological processes category involved “Phototransduction”, whereas “Hedgehog signaling pathway” was also observed to be enriched in the top 20 GO biological process **(figure 3B1)**. Out of the top 20 GO cellular component, we found 3 items “Photoreceptor outer segment”, “Photoreceptor disc membrane”, “Interphotoreceptor matrix” to be related to the retina **(figure 3B2)**. Of the top 20 GO molecular function, 3 terms were associated with axon development “regulation of axon extension”, “neural tube patterning” and “vagus nerve development”, besides GO associated with retinal function **(figure 3B3)**. Next, we focused our analysis on genes related to GO terms associated with retinal function and neuronal development **(figure 3C1-C5)**. We discovered a set of 22 downregulated genes that are associated with the GO biological process term “Phototransduction” **(Figure 3C1)**, and 7 genes linked to the GO molecular function term “Photoreceptor outer segment” that were downregulated **(Figure 3C3)**. We also detected 8 genes involved in the regulation of axon development with GO terms “regulation of axon extension” and “neuron projection” **(Figure 3C4)**.

**Figure 3:**
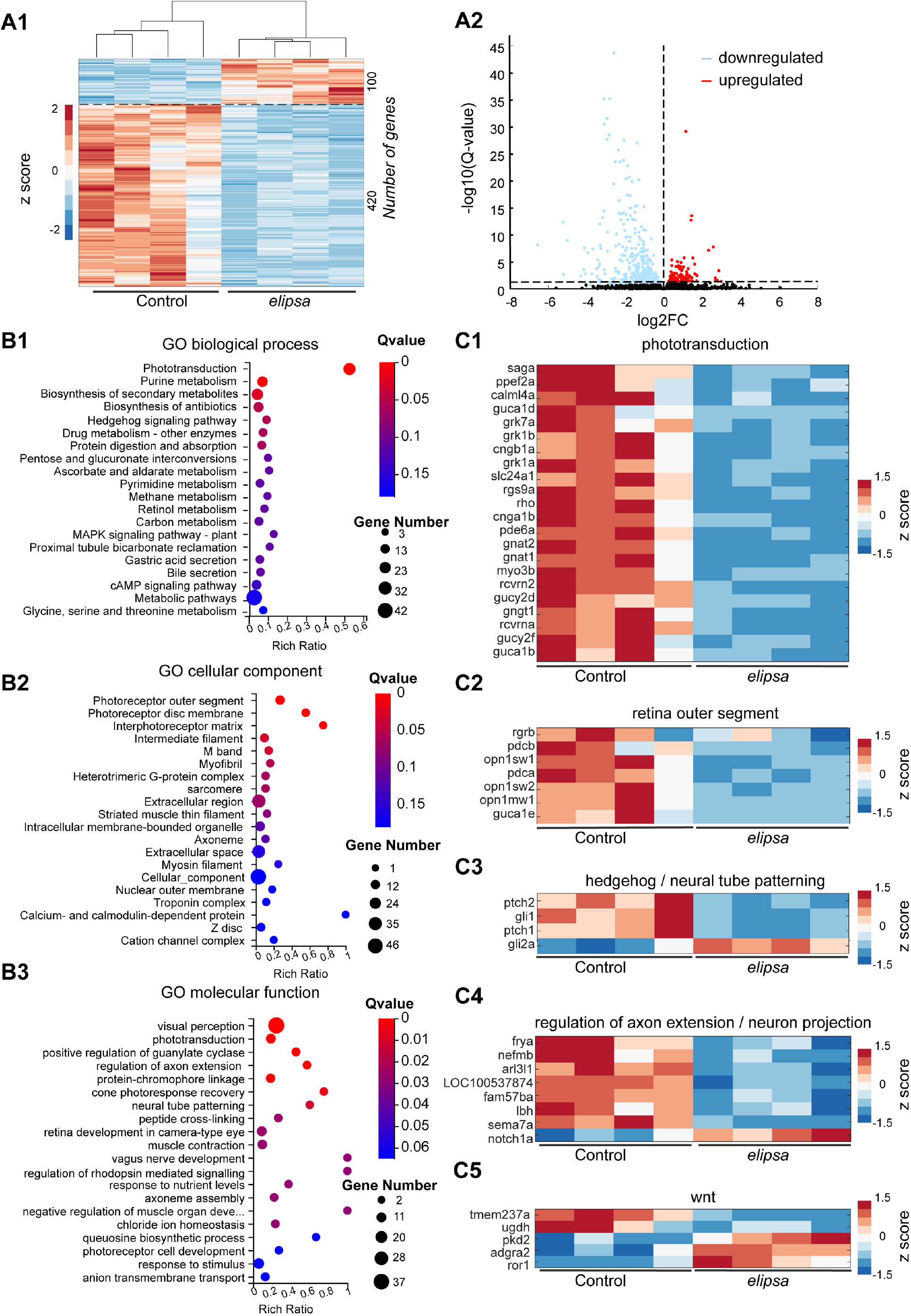
Transcriptomic analysis reveals altered expression of genes involved in phototransduction and brain development. **(A1)** Heatmap of expression of DEGs from control and *elipsa* 4dpf whole larvae (n =4 RNA preparation obtained from circa 30 control and *elipsa*) shown as z-score. Increased expression is represented with red and decreased expression with blue. A total of 520 (100 upregulated and 420 downregulated) differentially expressed mRNAs were identified. **(A2)** Volcano plot showing the DEGs in *elipsa* as compared with control. The horizontal dotted line indicates the Q-value (adjusted p-value) of 0.05. The vertical dotted line separate upregulated (red) and downregulated (blue) DEGs. **(B1-B3)** Top 20 Gene Ontology (GO) terms. **(B1)** Top 20 GO biological process **(B2)** Top 20 GO cellular component **(B3)** Top 20 GO molecular function. **(C1-C5)** Heat maps illustrate the expression levels of DEG associated with various GO terms **(C1)** “phototransduction” **(C2)** “retina outer segment” **(C3)** “hedgehog”/”neural tube patterning” **(C4)** “regulation of axon extension”/”neuron projection” and **(C5)** “Wnt” in control and *elipsa* larvae.

Furthermore, we investigated genes involved in cilia related signaling pathways such as the sonic hedgehog (shh) pathway (Bangs and Anderson, 2017; Wachten and Mick, 2021). Our analysis revealed 4 differentially regulated genes, including a downregulation of *ptch1, ptch2* and *gli1*, suggesting a dampened shh pathway **(Figure 3C2)**. Genes associated with neuronal development were mostly downregulated except for *notch1a*. Since Wnt signaling is essential for axon guidance and synapse development (Rosso et al., 2013), we also examined DEGs related to the Wnt signaling pathway and found that 5 genes showed differential regulation in the *elipsa* mutant **(Figure 3C5)**. Overall, our transcriptomics findings highlight a potential retinal defect in *elipsa* and the influence of cilia loss on specific signaling pathways, including shh and Wnt, within the brain.

### Cilia loss in *elipsa* larvae results in morphological and physiological defects in the photoreceptor layer

Since our transcriptomic analysis revealed downregulation of genes involved in phototransduction implying possible defects with the retina, we next sought to assess the morphology and physiological activity of the mutant’s retina. Notably, primary cilia were shown to play a crucial role in the formation and maintenance of photoreceptor outer segments in the vertebrate retina (Bachmann-Gagescu and Neuhauss, 2019). In line with this, prior work in the *elipsa* mutant identified a progressive degeneration of the photoreceptor outer segments at 5 dpf, with no morphological difference at 3 dpf (Bahadori et al., 2003). Here, we aimed to analyze the retinal phenotype at 4 dpf when mutant larvae remain relatively healthy beside their curved body axis. To this end we prepared retinal cryosections of 4 dpf control and *elipsa* larvae **(figure 4A1, B1)** and stained them using the lipophilic DiI dye. In comparison to the control larvae which had intact elongated outer segments **(figure 4A2)**, the outer segments of the *elipsa* larvae were shortened **(figure 4B2)**. Besides the photoreceptors, we did not observe major defects in the layering of the retina in both groups. Yet, we identified abnormal axonal projections from the retinal ganglion cells to the tectum in larvae with impaired cilia upon DiI injections **(figure 4C1-C2)**. Although all *elipsa* mutants displayed correct contralateral projections, the retinal ganglion cells commonly had circumvoluted tracts prior to the optic chiasm **(figure 4C2)**. Next, we investigated visual function by electroretinography (ERG) in 4 dpf larvae **(figure 4D1-D2)**, using a train of 1-second-long blue light stimuli.

**Figure 4:**
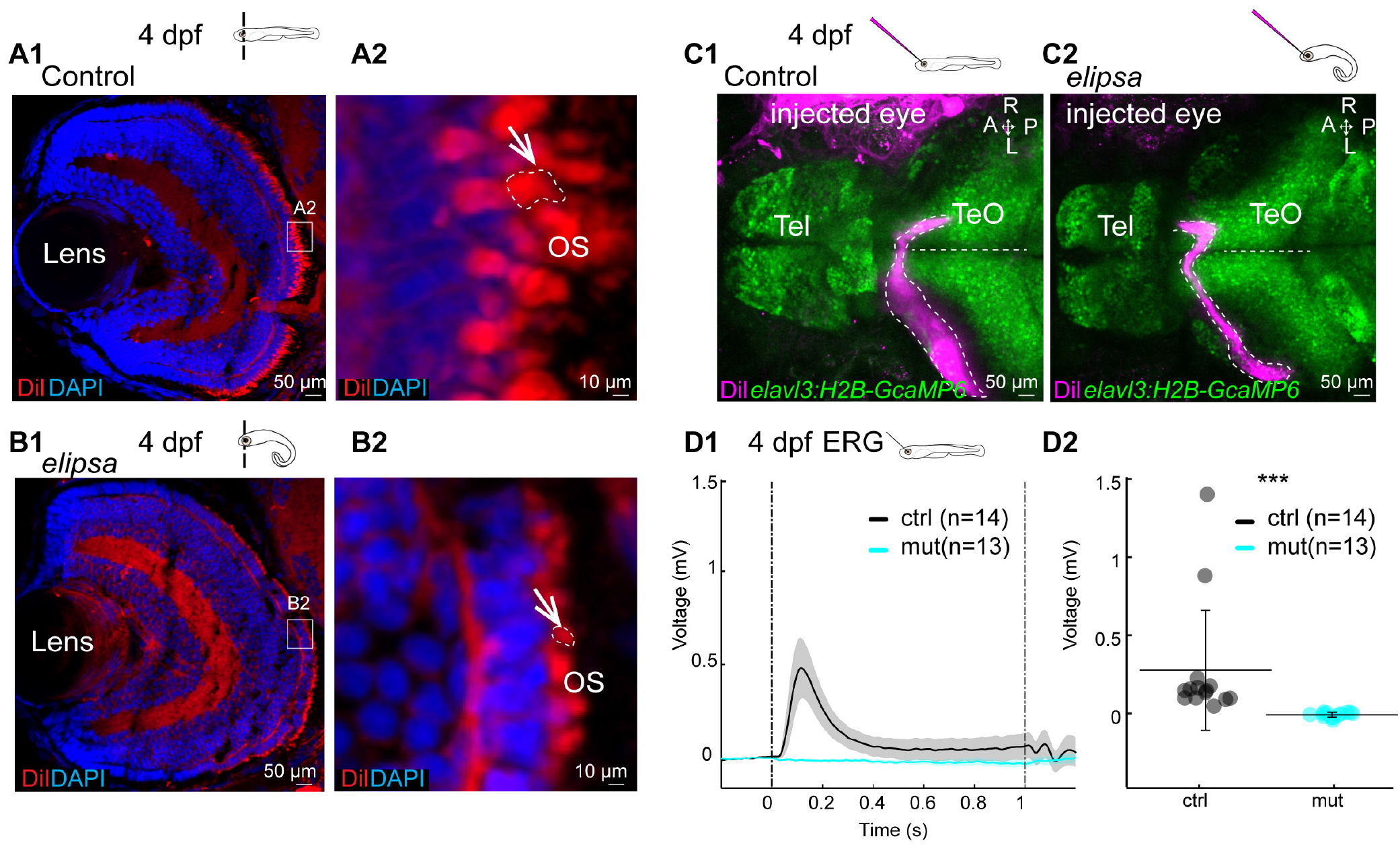
*elipsa* mutants display morphological defects of the photoreceptor outer segments and no retinal electrical activity. **(A1-B2)** DiI staining of 4 dpf retina cryosection to stain the outer segments. Section through the whole retina or the photoreceptor layer of a representative control **(A1-A2)** and *elipsa* **(B1-B2)**. Shortened outer segments (OS) are indicated using arrow and dashed lines. n =10 controls and 12 mutants. **(C1-C2)** DiI injection into the eye at 4dpf to stain the axonal connections (represented in dotted lines) which cross the midline and innervate the contralateral optic tectum. Here is shown a representative control **(C1)** and *elipsa* **(C2)**. Note that in the *elipsa* mutant the axonal tract goes posterior prior to returning to the optic chiasm. n =9 controls and 8 mutants. **(D1-D2)** Electroretinography (ERG) recordings in a 4 dpf retina. **D1** Average response of electrical activity (+/- standard error of the mean as the shaded region) to 1 second light stimulation for control (black) and mutant (cyan). **D2** Average electrical responses for the 200 msec following the light ON stimulus for all control fish (black) and *elipsa* mutant (cyan) showing no activity in the mutant. ***: p < 0.001 according to Wilcoxon Rank Sum test. Mean+/- standard deviation is indicated on scatter *plots*. n =14 controls and 13 mutants. Tel, Telencephalon; Teo, Optic Tectum.

We observed a complete absence of ERG signals in all analyzed *elipsa* as compared to controls. Taken together our results identified that cilia loss in the *elipsa* leads to morphological defects and loss of electrical activity in the retina already at 4 dpf.

### Reduced photic-induced neural activity in all brain regions of cilia-deficient larvae

To identify the impact of cilia loss on neuronal activity, we performed volumetric two-photon calcium imaging in 4 dpf *elipsa* larvae expressing the nuclear calcium indicator H2B-GCaMP6s in all differentiated neurons **(figure 5A1)** (Vladimirov et al., 2014). We performed 40 minute-long recordings consisting of spontaneous activity, followed by a series of photic stimulations that measure visual responses and can trigger seizure-like activity (Myren-Svelstad et al., 2022). We recorded 8 planes, used five light flashes of one minute each and collected data from four specific brain areas: the telencephalon, optic tectum/ thalamus, habenula, and hindbrain **(figure 5A2)**. Upon sorting cells based on k-means clustering, we observed distinct clusters of neurons exhibiting either enhanced, reduced or unchanged activity during photic stimulations, as shown by the representative examples **(figure 5A3-A4)**. To quantify neuronal response, we analyzed the neuronal activity during the light ON and OFF responses separately. We first examined the average activity of all neurons and found significantly reduced response for both ON **(figure 5B1-B5)** and OFF **(figure 5C1-C5)** conditions. In the controls, the optic tectum and hindbrain displayed the strongest responses to light flashes, while the telencephalon and habenula displayed comparatively minimal responses **(figure 5B2, C2)**. By computing the response amplitude for all cells in both light ON and OFF conditions **(figure 5B3 and C3)**, we observed reduced neuronal activity in most brain areas of the *elipsa* mutants. Notably, the tectum and hindbrain neurons exhibited reduced amplitudes for the light ON response condition **(figure 5B3)**, while all brain regions displayed lower amplitudes for the OFF condition **(figure 5C3)**. We then identified whether cells were generally less active or whether there were fewer responding cells. To determine the responding cells, we analyzed 10 seconds after the light stimulus and categorized cells with a mean amplitude greater than 2 standard deviations of the baseline as responding (as described in the methods). Our findings revealed a significant reduction in the number of activated cells across all brain regions for both light ON **(figure 5B4)** and OFF conditions **(figure 5C4)**. Moreover, we noted significant decrease in the number of inhibited cells in the telencephalon and optic tectum for the light ON condition **(figure 5B5)** and only in the optic tectum for the light OFF condition **(figure 5C5)**.

**Figure 5:**
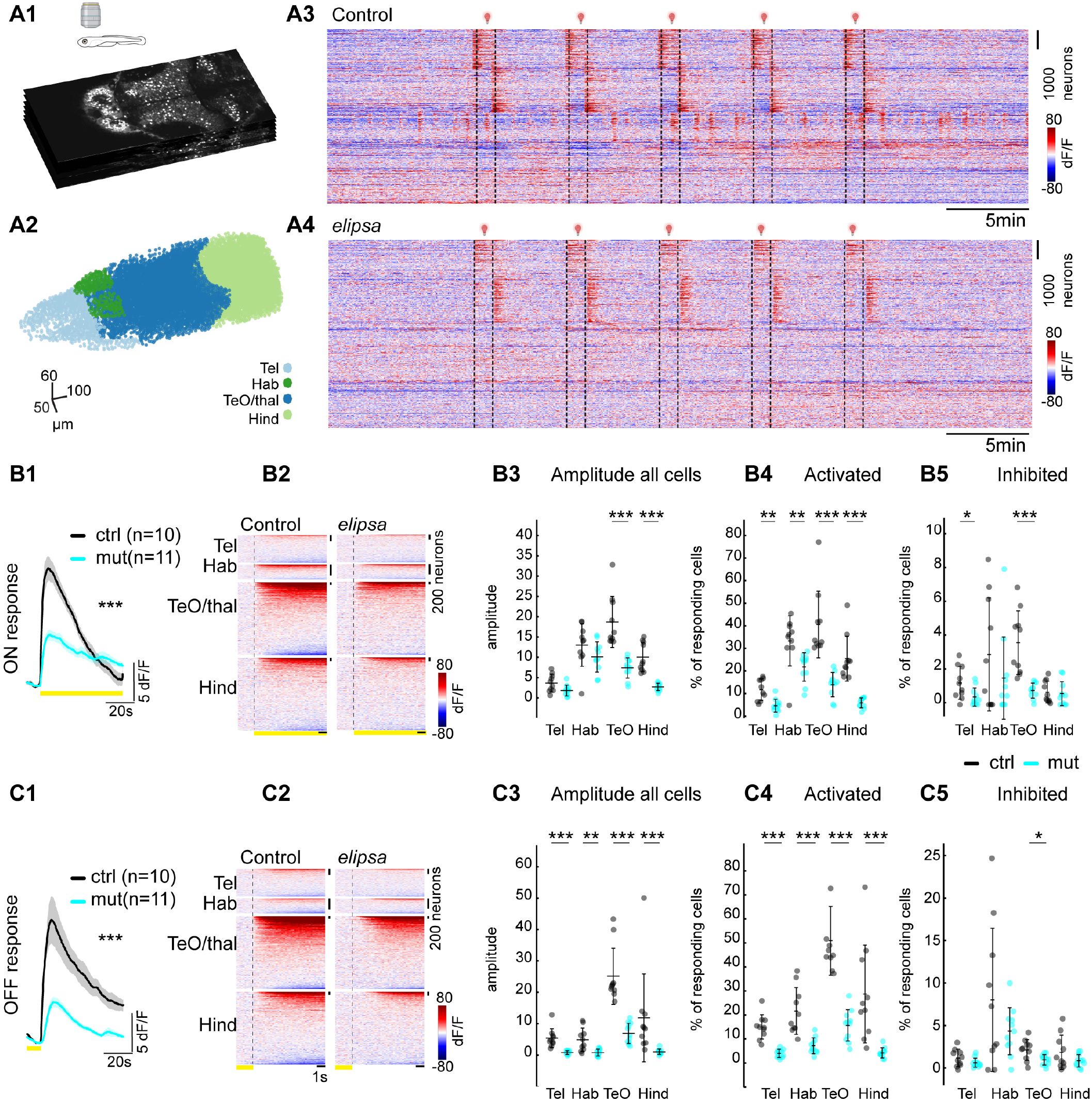
Reduced photic-induced neural activity in the brain of *elipsa* mutants. **A1** Optical sections of multiplane recording of a transgenic zebrafish larva expressing nuclear GCaMP6s in all neurons (*Tg(elavl3:H2B-GcaMP6s)*^*jf7Tg*^). **A2** Segmented nuclei from different brain regions, which were identified based on anatomical landmarks, are color coded. **A3-A4** Neuronal activity represented as change of fluorescence (dF/F) in one representative control and mutant larvae. Traces were sorted based on their activity using k-means clustering (warm color represents higher calcium signals). **B1, C1** Average response of neuronal activity to light stimulation for control (black) and mutant (cyan) for ON **(B1)** and OFF **(C1)** response conditions (+/- standard error of mean as the shaded region trace). **B2, C2** Activity of neurons per brain regions during ON **(B2)** and OFF response **(C2)** for representative examples (warm color represent higher calcium signals). **B3, C3** Average amplitude of all cells during ON **(B3)** and OFF **(C3)** response for control (black) and mutant (cyan), amplitude was significantly reduced in TeO and Hind regions for ON response and all brain regions for OFF response. **B4, C4** and **B5, C5** % of cells that are activated during ON **(B4)** and OFF response **(C5)** and inhibited during ON **(B5)** and OFF response **(C5)**. *: p <0.05, **: p <0.01, ***: p <0.001 according to to Wilcoxon Rank Sum test. Mean+/- standard deviation is indicated on scatter plots. n =10 controls and 11 mutants, control (black) and mutant (cyan). Tel, Telencephalon; TeO, Optic Tectum; BS, Brainstem; Hind, Hindbrain.

We previously identified that photic stimulation serves as a good strategy to detect hyperexcitability and trigger seizures in epilepsy models (Myren-Svelstad et al., 2022). We did not observed any epileptic seizures in the 11 mutant animals analyzed, suggesting that ciliary loss does not lead to the occurrence of seizure in the *elipsa* mutants at 4 dpf.

Taken together, our results using photic stimulation show that mutant animals exhibit reduced light-evoked neuronal activity in the entire brain, which aligns well with their retinal dysfunction. Interestingly, we measured residual neuronal activity throughout the brain of *elipsa* mutants despite the absence of ERG responses, suggesting that non retinal light-sensitive pathways remain at least partially active upon ciliary loss.

### Reduced spontaneous activity upon loss of cilia in *elipsa* mutant larvae

We next sought to investigate the impact of cilia loss on spontaneous brain activity. To this end, we quantified ongoing brain activity in control and *elipsa*. As shown in a representative example where neuronal activity was sorted based on their similarities using the rastermap (Carsen et al., 2023), *elipsa* mutants showed generally less activity in all brain regions **(figure 6A1-A4)**. Next, we measured the activity of cells by quantifying the percentage of time they spent above a threshold (see Methods) and generated an average cumulative frequency distribution for each brain regions. Our analysis showed that significantly fewer cells were highly active (active more than 50% of the time) in all brain regions in mutant animals **(figure 6B1-B4) (figure S3A1-D2)**. Concurrently, more cells were inactive (active less than 10% of the time) in the mutant **(figure 6B1-B4) (figure S3A1-D2)**. Furthermore, we investigated whether cells were differently correlated with each other. To this end we calculated the mean correlation versus distance for cells within distinct brain regions (Bartoszek et al., 2021; Diaz Verdugo et al., 2019; Fore et al., 2020). Our analysis revealed significantly less positive and negative correlation between nearby cells in the *elipsa* mutants **(figure 6C1-C4) (figure S3E1-H2)**. Altogether our results reveal that in absence of cilia neurons are less active and less correlated with each other, thereby emphasizing an important role of cilia in establishing functional neural networks in the zebrafish brain.

**Figure 6:**
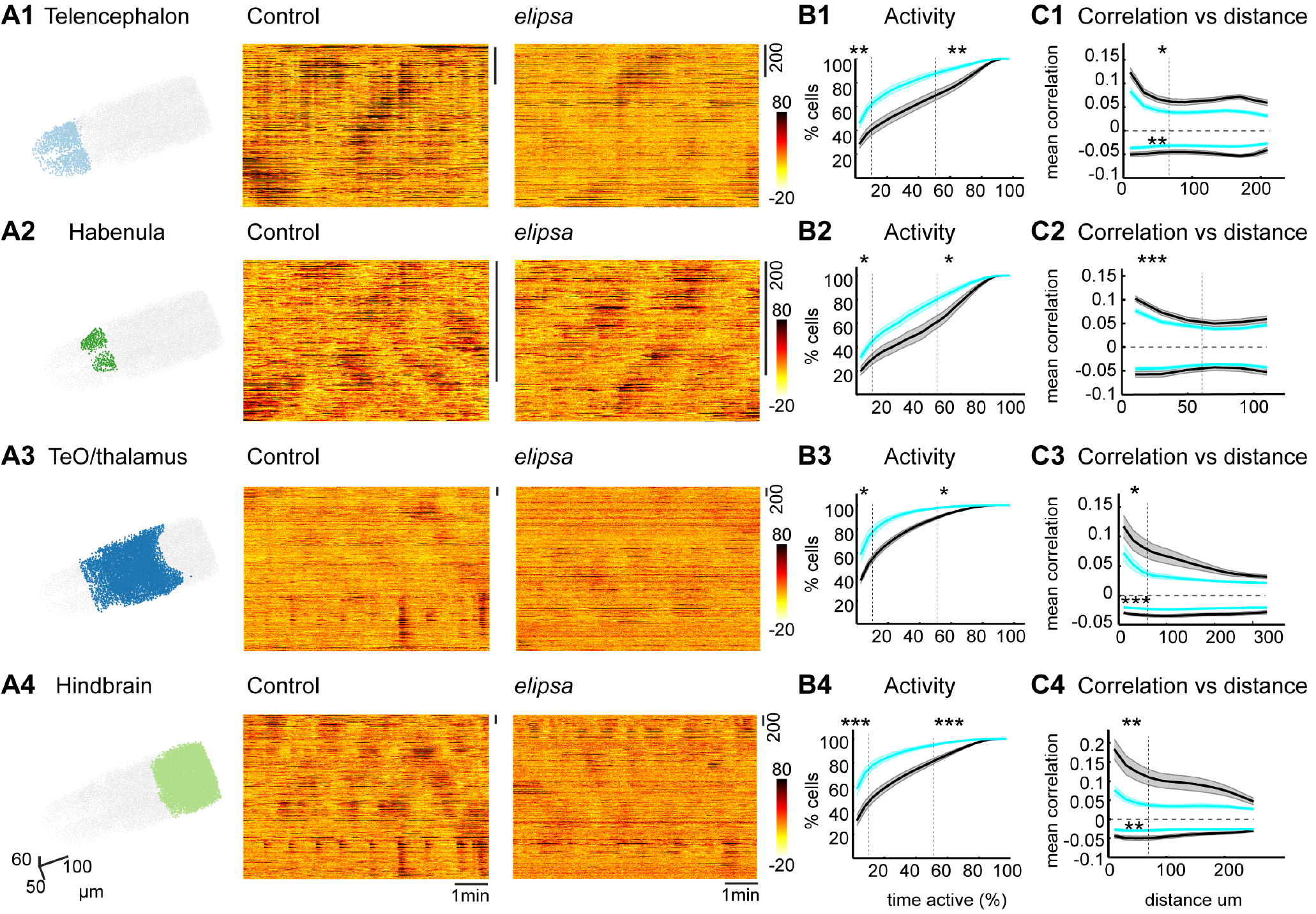
Reduced ongoing spontaneous activity and correlation in the *elipsa* mutants. **(A1-A4)** Three-dimensional representation of segmented neurons and their activity for one representative control and *elipsa* for the Telencephalon **(A1)**, Habenula **(A2)**, TeO/thalamus **(A3)** and Hindbrain **(A4)**. Dark color represents higher calcium signals. **(B1-B4)** Average cumulative frequency distribution graphs showing the percentage of activity of neurons in Telencephalon **(B1)**, Habenula **(B2)**, TeO/thalamus **(B3)** and Hindbrain **(B4)**, for control (black) and *elipsa* (cyan). (+/- standard error of mean as the shaded area). Dotted lines indicate activity below 10% and above 50%. **(C1-C4)** Mean correlation versus distance graphs showing positive and negative correlation between cells in Telencephalon **(C1)**, Habenula **(C2)**, TeO/thalamus **(C3)** and Hindbrain **(C4)**, for control (black) and mutant (cyan). Significance tests for the correlation were computed for cells located within 60 μm indicated by the dotted line on the X-axis. *: p <0.05, **: p <0.01, ***: p <0.001 according to Wilcoxon Rank Sum test . n =10 controls and 11 mutants.

## DISCUSSION

Cilia have been associated with a wide variety of functions in the developing brain, from neuronal proliferation and differentiation to axonal pathfinding and neuromodulation. Yet, how alterations in cilia-related neurodevelopmental processes impact brain activity have remained understudied. Moreover, despite zebrafish being a very common and powerful model to study cilia dysfunction, there has been no report on the function of cilia in the zebrafish brain. In our study, we took advantage of the small size of the zebrafish brain and its genetic toolbox to show that ciliary loss after neurulation alters brain development and the establishment of functional neural circuits using whole brain calcium imaging.

In our study, we chose specifically a zebrafish mutant line, *elipsa/traf3ip1* encoding for ift54, which has been studied earlier in the context of the retina, auditory hair cells, olfactory sensory neurons, spinal canal, brain ventricles and pronephros (Bahadori et al., 2003; Cantaut-Belarif et al., 2018; Doerre and Malicki, 2002; Olstad et al., 2019; Omori et al., 2008). In agreement with these prior works, we identified complete loss of cilia in the brain of *elipsa* mutants at stages following neurulation. Notably, we did not observe obvious ciliary defects in earlier developmental stages when the neural keel and Kupffer’s vesicle are established. The presence of cilia in the mutants at the 10 somites stage is most probably due to the maternal contribution of transcripts. Since we discovered that the mutant embryos start to lose their cilia between 13 hpf and 30 hpf, our observations indicate that ciliogenesis initially takes place but is subsequently lost during brain development and explain the absence of neural tube patterning defects in the *elipsa* mutant brains.

To date, cilia defects were mainly shown to lead to brain malformations in rodent models and human ciliopathy patients. These include abnormalities in the organization and connectivity of neurons and altered neuronal migration in various brain structures including the cortex, cerebellum, and hippocampus (Liu et al., 2021; Park et al., 2019; Suciu and Caspary, 2021; Youn and Han, 2018). In our study, we found that impairment of ciliogenesis had significant consequences for brain development also in zebrafish. We observed not only size differences for the telencephalon, optic tectum, and hindbrain but also malformations in the cerebellum. These findings align well with the phenotype of patients affected by Joubert’s syndrome (Bachmann-Gagescu et al., 2020; Joubert et al., 1969) which displays absence or underdevelopment of the cerebellar vermis as well as a malformed brain stem (Bashford and Subramanian, 2019; Brancati et al., 2010; Damerla et al., 2015; Ferland et al., 2004; Rachel et al., 2015; Rusterholz et al., 2022). Of note, cerebellar defects have only been described in a single zebrafish mutant so far, carrying a loss of function allele in the Joubert gene *arl13b* (Rusterholz et al., 2022; Zhu et al., 2020), and have been associated with reduced Wnt signaling. By identifying cerebellar malformation in the *elipsa* mutant, our findings are therefore opening new avenues for understanding cilia-related control of cerebellar development in Joubert’s syndrome using zebrafish as model system.

Since cilia serve as vital structures for the regulation of the cell cycle (Gabriel et al., 2016; Plotnikova et al., 2009; Tucker et al., 1979), we aimed to identify if the reduced brain size in our cilia mutant related to problems in cell proliferation. We did not observe any reduction in number of dividing cells but rather a significantly higher number of dividing cells in one specific brain area, the optic tectum. Some neural progenitors in the optic tectum are of neuroepithelial characteristics in contrast to the radial glia of the telencephalon (Jurisch-Yaksi et al., 2020; Lindsey et al., 2018; Recher et al., 2013). Therefore, it is possible that distinct types of neural progenitors are differentially affected by ciliary dysfunction and call for future work on identifying the molecular mechanisms underlying these differences.

Our transcriptomic analysis mainly uncovered genes linked to phototransduction and the formation of the outer segments of the retina. These results corroborated well with the photoreceptor abnormalities that we characterized, including a significant reduction in the size of outer segments and loss of retinal electrical activity. Our findings are consistent with previous work (Bahadori et al., 2003), which described the impact of the *elipsa* mutation on the structure and function of the retina at 5 dpf. It is important to note that despite observing a fully penetrant retinal dysfunction at 5 dpf, this study did not report any apparent morphological abnormalities at 3 dpf (Bahadori et al., 2003), supporting the fact that cilia-related retinal defects are usually progressive and worsens over time. We now provide further evidence that the retina is already affected at 4 dpf, a developmental stage when larvae remain relatively healthy to be analyzed in depth for neuronal function.

We also uncovered genes related to hedgehog and Wnt signaling pathways, neural patterning, regulation of axonal extension and vagus nerve development. Hedgehog signaling has emerged as a pivotal regulatory pathway governing the growth and patterning of the cerebellum. (Dahmane and Ruiz i Altaba, 1999; Lee et al., 2010; Wallace, 1999; Wang et al., 2022; Wechsler-Reya and Scott, 1999). This suggests that dampening of shh signaling upon cilia loss could contribute to the cerebellar malformations of the *elipsa* mutants. Of note, our transcriptomics analysis was performed on whole larvae and may have missed genes that are expressed at low levels or in restricted cell populations in the brain.

To analyze the impact of ciliary loss on brain physiology, we used volumetric two-photon calcium imaging and measured spontaneous and light-induced neuronal activity. In zebrafish, light stimulation elicits neuronal responses in various brain regions. These include the visual center known as the optic tectum (Robles et al., 2014), the thalamus (Mueller, 2012), which receive direct input from the retinal ganglion cells, and their downstream projections to the habenula (Dreosti et al., 2014; Zhang et al., 2017), telencephalon and hindbrain (Mueller, 2012). In line with retinal defects, we observed a significant decrease in the light response across all brain regions in *elipsa* larvae. Notably, we did not observe a complete loss of light-induced neuronal activity in the *elipsa* mutants. This could be related to a potential residual photoreceptor activity in the mutant, which were too small to be detected by ERG, or non-retinal light responses. The pineal gland, a ciliated organ located dorsally above the telencephalon (Laurà et al., 2012; Vigh et al., 2002), may be involved, although one could expect that cilia loss would affect the pineal responses too. Alternatively, it has been reported that specific population of neurons express light-responsive opsins, which regulate animal behavior (Dekens et al., 2022; Fernandes et al., 2012; Fontinha et al., 2021). These neurons may therefore be involved in the light responses that remained in the *elipsa* mutants. One prior report identified that *elipsa* morphants (Lepanto et al., 2016) displayed altered proliferation of retinal progenitors and survival of RGCs. Therefore, it is possible that the reduced light-evoked neuronal activity in the *elipsa* mutants stems from an accumulation of developmental defects, loss of outer segments and reduced photoreceptor activity.

Our investigation into spontaneous activity (Bartoszek et al., 2021; Fore et al., 2020; Jetti et al., 2014) revealed notable differences in the *elipsa* mutants including significant reduction in the number of active cells and correlation between cells. These differences can stem from the neurodevelopmental defects that we have observed. Since spontaneous activity can play a role in maturation of functional circuits, and refinement of axonal projections (Choi et al., 2021; Kirkby et al., 2013), it is possible that the reduced spontaneous activity further exacerbates defects in neuronal development and gene expression. Altered spontaneous activity in the brain is observed in various conditions, including neurodevelopmental disorders, neurodegenerative diseases, or brain injuries, which usually manifest with intellectual disabilities or cognitive impairments (Liu et al., 2022; Uddin, 2020). Notably, ciliary dysfunction in Joubert and Bardet-Biedl syndromes is commonly associated with intellectual disabilities (Bachmann-Gagescu et al., 2015; Forsyth and Gunay-Aygun, 1993; Forsythe and Beales, 2013; Parisi and Glass, 1993), suggesting that it may originate from altered spontaneous activity. Overall, while the precise link between cilia and spontaneous activity in the brain is not fully established, future experiments manipulating cilia dynamics in specific neurons will be able to uncover the specific molecular mechanisms controlling cilia-dependent brain activity.

Ciliopathy syndromes including Joubert and Bardet-Biedl syndrome are associated with epilepsy (Bachmann-Gagescu et al., 2015; Forsyth and Gunay-Aygun, 1993; Forsythe and Beales, 2013), which is a neurological disorder characterized by recurrent seizures. The exact mechanisms underlying the association between ciliopathies and epilepsy are not fully understood and may vary depending on the specific syndrome. While our study revealed cilia-related neurodevelopmental defects, we did not observe any spontaneous or light-evoked epileptic seizures in the *elipsa* mutant larvae. This may be related to the analyzed developmental stage, 4 dpf, which may be too early to detect seizure-like activity. Indeed, while some zebrafish epilepsy models, including the mutant carrying a loss-of-function mutation in the glutamate transporter *eaat2a*, were shown to already display seizures at 5 dpf (Hotz et al., 2022), other models develop seizure only at the juvenile stages like the GABA A receptor mutants (Samarut et al., 2018; Yaksi et al., 2021). Alternatively, one could argue that reduced retinal activity could prevent light-induced hyperexcitability in the *elipsa* mutant. Yet, the *eaat2a* mutants, which also show reduced retinal activity (Niklaus et al., 2017), display strong photic-induced epileptic seizures (Hotz et al., 2022; Myren-Svelstad et al., 2022).

In summary, our results demonstrate that loss of cilia post neurulation has impacts on the development of functional neural circuits and lead to altered neuronal activity in the larval zebrafish brain. We are confident that this study opens avenues to study the molecular mechanisms underlying cilia-related brain dysfunction in ciliopathy models and identify potential therapeutical targets.

## ACKNOWLEDGEMENTS

We thank Zhaoxia Sun for sharing the Arl13b antibody, our fish facility team for husbandry maintenance and technical support, and all members of the Jurisch-Yaksi and Yaksi laboratories for their feedback on this work and exchanging MATLAB codes. This work was supported by funding from an NTNU strategy grant (NJY) and The Research Council of Norway: RCN FRIPRO grant 314189 (NJY).

## AUTHOR CONTRIBUTIONS

Conceptualization: NJY, PPD; Methodology: PPD, IJ, AN, ATT, EY; Formal analysis: PPD, IJ, NJY, ATT; Investigation: PPD, IJ, AN, NJY; Resources: NJY, EY; Data curation: PPD, IY, NJY; Writing - original draft: PPD, NJY; Writing - review & editing: all authors; Visualization: PPD, NJY; Supervision: NJY, EY; Funding acquisition: NJY, EY.

## DECLARATION OF INTERESTS

The authors declare that there is no conflict of interest

## MATERIAL AND METHODS

### Lead contact

Further information and requests for resources and reagents should be directed to and will be fulfilled by the lead contact, Nathalie Jurisch-Yaksi.

### Zebrafish maintenance and strains

The animal facilities and maintenance of the zebrafish, *Danio rerio*, were approved by the NFSA (Norwegian Food Safety Authority). All the procedures were performed on zebrafish were in accordance with the European Communities Council Directive, the Norwegian Food Safety Authorities. The larval and adult zebrafish were reared according to standard procedures of husbandry at 28.5 °C, in 3.5 L tanks in a Techniplast Zebtech Multilinking system at constant pH 7 and 700 μSiemens, at a 14:10 hr light/dark cycle to mimic optimal natural breeding conditions. Larvae were maintained in egg water (1.2 g marine salt and 0.1% methylene blue in 20 L RO water) from fertilization to 3 dpf and subsequently in AFW (1.2 g marine salt in 20L RO water). For our experiments, the following fish line were used *elipsa/traf3ip1*^*tp49d*^ (received from J Malicki, University of Sheffield) and *Tg(elavl3:H2B-GcaMP6)*^*jf7Tg*^. Two photon calcium experiments were performed with larvae obtained from incrossing heterozygous *elipsa*^*+/-*^;*Tg(elavl3:H2B-GCaMP6s)* ^jf7Tg^ adult animals. For immunostaining and RNA sequencing, larvae were obtained by crossing heterozygous *elipsa*^*+/-*^ animals. Controls were either wild-type or *elipsa*^+/-^ obtained from the same breeding.

### Genotyping

For genotyping, the samples were subjected to gDNA isolation using 100 μL PCR lysis buffer (containing 1M tris pH-7-9, 0.5 M EDTA, Triton-100 and Proteinase K 0.1mg/ml) overnight at 50 °C. To stop the reaction the samples were heated to 95°C for 10 minutes. The samples were then centrifuged at 13000 rpm for 2 minutes. The supernatant containing gDNA was used for further KASP assays-based analysis. The samples were first diluted (1:2) with water. Further 3 μL of diluted sample was used for performing the KASP assay according to the manufacturer guidelines. The master mix contained 5 μL mastermix, 0.14 μL assay mix and 1.86 μL milliQ water per sample well.

### Antibody staining and confocal imaging Immunostaining of the brain with cilia specific antibodies

Larvae were euthanized and fixed in a solution containing 4 % paraformaldehyde solution (PFA), 1 % DMSO and 0.3 % TritonX-100 in PBS (0.3 % PBSTx) for 2 hours at room temperature or 4 °C overnight. The larvae were washed with 0.3 % PBSTx after fixing to remove any traces of the fixing solution. For permeabilization, samples were incubated for 10 minutes at -20 °C with acetone. Subsequently, samples were washed with 0.3 % PBSTx (3x10 min) and blocked in 0.1 % BSA made in 0.3 % PBSTx at room temperature. Samples were incubated with the primary antibody overnight at 4 °C. The antibodies used were mouse glutamylated tubulin (GT335, 1:400, for staining motile cilia), rabbit Arl13b antibody (1:200, for staining all cilia), or rabbit p-Histone H3 Antibody (1:500, for staining diving cells). On the second day samples were washed (0.3 % PBSTx, 3x1 hour) and subsequently incubated with the secondary antibody (Alexa-labelled GAM488 plus, or GAR555 plus Thermo Scientific, 1:1,000) and 0.1 % DAPI overnight at 4 °C. On the third day after incubation with the secondary antibody the larvae were washed (0.3 % PBSTx, 3x1 hour) and transferred to a series of increasing glycerol (made in PBS) concentrations (25 %, 50 % and 75 %). After staining the larvae were stored in 75 % glycerol at 4 °C and imaged using a Zeiss Examiner Z1 confocal microscope with a 20x plan NA 0.8 objective. Multiple images were stitched using Fiji (Preibisch et al., 2009). For detailed protocol refer (D’Gama and Jurisch-Yaksi, 2023).

### Quantification of pH3+ cells and measuring brain morphology

After image acquisition, the pH3+ cells were counted using the cell counter function in fiji (https://imagej.nih.gov/ij/plugins/cell-counter.html). The brain morphology measurements were done on Z stack using the straight-line tool in Fiji.

### Cryo-sectioning and staining of the retina

The protocol for cryo-sectioning was adapted from (Masek et al., 2023). Zebrafish larvae were euthanized using ice water (4°C). The larvae were then fixed using 4% PFA in 0.3% triton-100 PBS overnight at 4°C. After fixing the samples were washed 3x1hour in PBS. The fish were embedded before cryoprotection, using a solution of 1.5% agarose and 5% sucrose solution in RO. The larvae were transferred to a cryomold, and the embedding solution was added to this mold. The fish were positioned using a pick and the solution was allowed to cool. After cooling, the mold was cut into tiny blocks (cut the corner to indicate orientation). The blocks were then transferred to storage solution (30% Sucrose in PBS) and stored in the fridge overnight at 4°C until the samples sank to the bottom of the tube. The next day, the blocks containing the samples were snap frozen with liquid nitrogen using a metal bowl with 2-methylbutane that is positioned in a Styrofoam container. Frozen samples were stored at -20°C until further use. Using a cryostat, sections of 10 μm thickness were cut at -30°C blade and chamber temperature. The sections were placed on super frost slides and stored at -20°C until further use. The cryosection slides were thawed at RT for 30 minutes. The slides were washed with PBS without detergent for 4x5 minutes to remove any traces of freezing medium. The outer segments were stained using DiI stain (5mg/ml diluted 1:100-1:200 in PBS) for 20 minutes at RT, followed by 3x5 minute washes with 1X PBS. The slides were then dried and mounted with prolong gold. The slides were left overnight to dry before imaging.

### DiI-based tracing of retinal projection

4 dpf larvae were euthanized and then fixed with 4% PFA in PBS at 4 dpf O/N. Following 3 x1 hour washes with PBS, fish were immobilized with 1.5% LMP agarose in RO water. Injections into the eye were performed using an Eppendorf Femtojet 4i pressure injector and glass capillaries that were filled with 5 mg/ml DiI diluted in DMF. Following screening the larvae were imaged using a Zeiss Examiner Z1 confocal microscope with a 10x plan NA 0.45 objective.

### RNA sequencing and transcriptomic analysis

To isolate RNA for sequencing, 30 larvae at 4 dpf were collected in a 1.5ml tube and placed on ice. To lyse the samples, 500μL trizol was added and the euthanized larvae were homogenized through a 27-gauge needle until the mixture looked uniform. After adding another 500μL trizol, the samples were incubated for 5 minutes at room temperature. The larvae were then treated with 200μL chloroform, and the tube was rocked for 15secs to mix the contents. The tubes were incubated for 2 minutes at room temperature and then centrifuged for 15 minutes at 12000rpm at a temperature of 4°C. After centrifugation, the upper aqueous phase containing RNA was mixed with equal amounts of 100% ethanol and was then loaded onto an RNA spin column (Qiagen) and centrifuged for 30 seconds at 8000 rpm. The spin column was further incubated with 700μL of RW1 buffer and centrifuged for 30 seconds at 8000 rpm. The spin column tubes were then placed into a new collection tube and further treated to remove any DNA contamination by washing the tubes with 350μL of RW1 buffer followed by Dnase enzyme (Qiagen) in RDD buffer (10μL Dnase+ 70μL RDD buffer per tube) for 45 minutes at room temperature. After incubation, 350μL of RW1 buffer was added to the tubes and centrifuged for 15 seconds at 8000rpm. The tubes were then treated with 500μL RPE buffer and centrifuged for 30 seconds. This step was repeated twice, and the tubes were then centrifuged for 1 minute at 8000rpm to remove any residual buffer left in the column. For RNA extraction from the column, 30μL nuclease free water was added and incubated for 2 minutes. The tubes were then centrifuged for 1 minute at 8000 rpm to elute the RNA. The concentration of the extracted RNA was quantified using Nanodrop and the quality was analyzed by bioanalyzer. The samples were then sequenced by BGI’s DNBSEQTM Technology using the Dr Tom data visualization and analysis platform provided by BGI.

### Two photon calcium imaging

Two-photon calcium imaging was performed on 4 dpf *elipsa*;*Tg(elavl3:H2B-GCaMP6)* ^*jf7Tg*^. The larvae were paralyzed upon injection of α-bungarotoxin (Invitrogen BI601, 1 mg/ml) and embedded in 1.5% low melting point agarose in mounting chambers (Fluorodish, World Precision Instruments) using a plastic microcapillary tip, as described in (Reiten et al., 2017). After 10 min of agarose solidification, 750 μL of AFW was added on top of the agarose. The mounting chamber was then allowed to stabilize under the two-photon microscope before recording. The recordings were performed in a two-photon microscope (Scientifica) using a 16× water immersion objective (Nikon, numerical aperture 0.8, Long Working Distance 3.0, plan) and a Ti:Sapphire laser (MaiTai Spectra-Physics) tuned at 920 nm. Recordings of 1536 ×512 pixels and 8 planes were acquired at a frame rate of 30.85 Hz and a volume rate of 3.86 Hz. The total recording time was 40 minutes. Ongoing activity was recorded for 10 minutes in darkness, followed by 5 stimuli each of 60 seconds using a red light-emitting diode light (LZ1-00R105, LedEngin; 625 nm), at minute 10, 15, 20, 25 and 30 respectively. Animals without cerebral blood flow after the experiments were excluded from the analysis.

### Data analysis

Two-photon microscopy images were aligned using a previously reported algorithm (Fore et al., 2020; Reiten et al., 2017). The recordings were then screened manually to check for movement and Z drift. Unstable recordings were discarded from the analysis. All analyses were performed on MATLAB. Neurons were segmented using a pattern recognition algorithm adapted from (Ohki et al., 2005) with torus or ring shaped neuronal templates (Jetti et al., 2014). The identified neurons were then set apart into different brain regions based on another algorithm described in (Bartoszek et al., 2021) **(figure 5A2)**. Clustering of neurons was performed using k-means clustering algorithm based on their activity (Jetti et al., 2014). For detecting light responses, we used the 5 seconds of fluorescence preceding the stimuli as baseline and average of the 5 stimuli for each neuron. The 5s baseline was used to calculate dF/F. Responsive cells were selected based on their response during the 10 seconds following the light onset or offset. Cells were classified as responsive if their average activity during the 10 second window was greater than 2* standard deviation of the baseline fluorescence.

For measuring spontaneous activity, we excluded the first two minutes of the recordings (to avoid artefacts from the microscope laser turning on). We selected frames 460-2304 (a total of 7.5 minutes of ongoing activity). dF/F was calculated using, as baseline, the 8th percentile of a moving window as previously described (Fore et al., 2020; Romano et al., 2017).The data was resampled to 1fps using the resample function in MATLAB Calculations for the activity of cells were done on resampled data. A cell was considered active at a given second if it had an activity higher than a threshold of 4* 8th percentile. We then calculated the percentage of cells that are very active (active more than 50% of the time) or inactive (active less than 10% of the time) per brain region. The Pearson’s correlation versus distance between neurons was calculated up to 60μm per brain region **(figure S3)**.

### Electroretinography

Electroretinography (ERG) were performed in control or *elipsa* mutants at 4dpf. First, we selected healthy control or *elipsa* mutants at 4dpf and anesthetized the larvae by using MS222. We then placed the anesthetized larvae on a wet filter paper (VWR, 516-0848) on a FluoroDish (VWR, FD35PDL-100) and covered their trunk with paper towels to affix them. We used a sliver-coated wire as a reference electrode, which was chlorinated by immersing in sodium hypochlorite solution 10 minutes before the recordings. The reference electrode was positioned on the same bath as the larvae. Recording electrodes were pulled from glass capillaries (WPI, TW100F-4) to have a 15-30 μm opening and filled with artificial fish water (AFW) containing MS222. The electrode was placed on the cornea of the eye using a motorized micromanipulator (Scientifica). Before the recording, larvae were subjected to over 10 minutes dark adaption. The voltages were measured by a MultiClamp 700B device (Molecular Devices) with a 2kHz low-pass filter and were digitized at 10 kHz. A train of 10 light stimuli was initiated at the 5th second of the recording. The stimuli were produced by a blue LED using a pulse generator (Master-8, AMPI). Each stimulus (light intensity: 0.005-0.007 mW) lasted 1 sec with 5 seconds intervals between stimuli. The total time for each recording was 60 seconds. We performed minimum 5 recordings for each larva and selected the trial that shows the most representative responses for further analysis. We utilized custom MATLAB scripts for data acquisition and all subsequent analyses. To quantify the voltage responses, we used a 200 msec baseline before the light stimulus to normalize the traces and averaged the 10 light stimuli. We calculated the average voltage responses for all control and mutants from 0 to 200 msecs after the light ON stimuli, which show the strong b-wave representing the ON bipolar cell respons

### Quantification and statistical analysis

Statistical analysis was done using MATLAB. Wilcoxon rank-sum test was used for nonpaired analysis. Probability of p < 0.05 was considered statistically significant.

| REAGENT or RESOURCE | SOURCE | IDENTIFIER |
| --- | --- | --- |
| **Antibodies** |  |  |
| Mouse monoclonal glutamylated tubulin (GT335) | Adipogen | Cat#AG-20B-0020-C100;<br>RRID: AB_2490210 |
| Rabbit Arl13b antibody | Zhaoxia Sun Lab | (Duldulao et al., 2009) |
| Anti-acetylated tubulin antibody | Sigma | Cat#T7451;<br>RRID:AB_609894 |
| Rabbit p-Histone H3 Antibody | Santacruz | Cat#sc-8656-R;<br>RRID:AB_653256 |
| Alexa fluor plus Goat anti rabbit 555 | Thermo Fisher Scientific | Cat#A32732;<br>RRID:AB_2633281 |
| Alexa fluor plus Goat anti mouse 555 | Thermo Fisher Scientific | Cat#A32727,<br>RRID:AB_2633276 |
| Alexa fluor plus Goat anti mouse 647 | Thermo Fisher Scientific | Cat#A32728;<br>RRID:AB_2633277 |
| **Chemicals** |  |  |
| DAPI (4',6-Diamidino-2-Phenylindole, Dihydrochloride) | Thermo Fisher Scientific | Cat#D1306;<br>RRID:AB_2629482 |
| Microamp optical 96 well reaction plate (Applied biosystems) | Thermo Fisher Scientific | Cat# N8010560 |
| Phosphate buffered saline | Thermo Fisher | Cat#BR0014G |
| Triton X-100 | Merck | Cat# 1086031000 |
| Bovine Serum Albumin (BSA) | PanReac AppliedChem | Cat#A1391 |
| Dimethyl sulfoxide (DMSO) | Sigma | Cat#D8418 |
| Glycerol | VWR | Cat#24387.292 |
| Acetone | VWR | Cat#20066.296 |
| Formaldehyde solution (PFA) | Sigma | Cat#F8775-25 |
| MS-222 | Sigma | Cat#E10621-60G |
| Proteinase K from tritirachium album | Sigma | Cat#P2308-25MG |
| Tris | Sigma | Cat#252859-100G |
| Alpha-bungarotoxin | Invitrogen | Cat#B1601 |
| Agarose | Thermo Scientific | Cat#P17850 |
| Sucrose | Invitrogen | Cat# 15503022 |
| Dil stain (1,1'-Diocetadecyl-3,3',3'-Tetramethylindocarbocyanine Perchlorate ('Dil'; DiIC18(3))) | Invitrogen | Cat#D282 |
| ProLong Gold Antifade Mountant | Invitrogen | Cat#P36930 (10 ml) |
| Trizol | Invitrogen | Cat# 15596026 |
| Rneasy mini kit | Qiagen | Cat#74104 (50 reactions) |
| Rnase-Free Dnase Set | Qiagen | Cat#79254 (50 reactions) |
| Ethanol (molecular grade) | Merck | Cat#1.00983 |
| Ultra LMP agar | Invitrogen | Cat#16520-100 |
| Thin wall capillary 1 mm | World precision instruments | Cat#TW100F-4 (with filament) |
| Filter paper | VWR | Cat#516-0848 |
| Fluorodish | VWR | Cat# FD35PDL-100 |
| Glass capillary | WPI | Cat# TW100F-4 |
| **Experimental models: Zebrafish** |  |  |
| *Elipsa traf3ip1<sup>tp49d</sup>* | (Omori et al., 2008) | RRID:ZFIN_ZDB-ALT-980413-466 |
| *Tg(elavl3:H2B-GCaMP6s)* <sup>jf7Tg</sup> | (Vladimirov et al., 2014) | ZFIN: ZDB-ALT-150916-4 |

| Commercial Assays |  |  |
| --- | --- | --- |
| KASP Assay | LGC genomics | NA |
| Other |  |  |
| Pressure injector | Eppendorf | Femtojet 4i |
| Confocal microscope | Zeiss | Examiner Z1; Olympus Fluoview |
| Two photon microscope | Scientifica | NA |
| Step One Real Time PCR system | Thermofisher | Cat#4376357 |
| Sutter Laser puller | Sutter | Model P-2000 |
| Cryostat NX 70 | Thermo Scientific | NA |

## Notes

### Competing Interest Statement

The authors have declared no competing interest.

## References

1. Andreu-Cervera, A., Catala, M. and Schneider-Maunoury, S. (2021). Cilia, ciliopathies and hedgehog-related forebrain developmental disorders. Neurobiology of Disease 150, 105236.

2. Bachmann-Gagescu, R., Dempsey, J. C., Bulgheroni, S., Chen, M. L., D’Arrigo, S., Glass, I. A., Heller, T., Héon, E., Hildebrandt, F., Joshi, N., et al. (2020). Healthcare recommendations for Joubert syndrome. American Journal of Medical Genetics Part A 182, 229–249.

3. Bachmann-Gagescu, R., Dempsey, J. C., Phelps, I. G., O’Roak, B. J., Knutzen, D. M., Rue, T. C., Ishak, G. E., Isabella, C. R., Gorden, N., Adkins, J., et al. (2015). Joubert syndrome: a model for untangling recessive disorders with extreme genetic heterogeneity. J Med Genet 52, 514–522.

4. Bachmann-Gagescu, R. and Neuhauss, S. C. F. (2019). The photoreceptor cilium and its diseases. Current Opinion in Genetics & Development 56, 22–33.

5. Bahadori, R., Huber, M., Rinner, O., Seeliger, M. W., Geiger-Rudolph, S., Geisler, R. and Neuhauss, S. C. (2003). Retinal function and morphology in two zebrafish models of oculo-renal syndromes. Eur J Neurosci 18, 1377–1386.

6. Bangs, F. and Anderson, K. V. (2017). Primary cilia and mammalian hedgehog signaling. Cold Spring Harbor perspectives in biology 9, a028175.

7. Bartoszek, E. M., Ostenrath, A. M., Jetti, S. K., Serneels, B., Mutlu, A. K., Chau, K. T. P. and Yaksi, E. (2021). Ongoing habenular activity is driven by forebrain networks and modulated by olfactory stimuli. Curr Biol 31, 3861–3874.e3863.

8. Bashford, A. L. and Subramanian, V. (2019). Mice with a conditional deletion of Talpid3 (KIAA0586) - a model for Joubert syndrome. J Pathol 248, 396–408.

9. Bear, R. M. and Caspary, T. Uncovering cilia function in glial development. Annals of Human Genetics n/a.

10. Berbari, N. F., Lewis, J. S., Bishop, G. A., Askwith, C. C. and Mykytyn, K. (2008). Bardet-Biedl syndrome proteins are required for the localization of G protein-coupled receptors to primary cilia. Proc Natl Acad Sci U S A 105, 4242–4246.

11. Bergboer, J. G. M., Wyatt, C., Austin-Tse, C., Yaksi, E. and Drummond, I. A. (2018). Assaying sensory ciliopathies using calcium biosensor expression in zebrafish ciliated olfactory neurons. Cilia 7, 2.

12. Brancati, F., Dallapiccola, B. and Valente, E. M. (2010). Joubert Syndrome and related disorders. Orphanet J Rare Dis 5, 20.

13. Bujakowska, K. M., Liu, Q. and Pierce, E. A. (2017). Photoreceptor Cilia and Retinal Ciliopathies. Cold Spring Harbor Perspectives in Biology 9.

14. Cantaut-Belarif, Y., Sternberg, J. R., Thouvenin, O., Wyart, C. and Bardet, P. L. (2018). The Reissner Fiber in the Cerebrospinal Fluid Controls Morphogenesis of the Body Axis. Curr Biol 28, 2479–2486.e2474.

15. Carsen, S., Lin, Z., Atika, S., Fengtong, D., Maria, K. and Marius, P. (2023). Rastermap: a discovery method for neural population recordings. bioRxiv, 2023.2007.2025.550571.

16. Choi, B. J., Chen, Y. D. and Desplan, C. (2021). Building a circuit through correlated spontaneous neuronal activity in the developing vertebrate and invertebrate visual systems. Genes Dev 35, 677–691.

17. D’Gama, P. P. and Jurisch-Yaksi, N. (2023). Methods to study motile ciliated cell types in the zebrafish brain. In Methods in Cell Biology: Academic Press.

18. D’Gama, P. P., Qiu, T., Cosacak, M. I., Rayamajhi, D., Konac, A., Hansen, J. N., Ringers, C., Acuña-Hinrichsen, F., Hui, S. P., Olstad, E. W., et al. (2021). Diversity and function of motile ciliated cell types within ependymal lineages of the zebrafish brain. Cell Rep 37, 109775.

19. Dahmane, N. and Ruiz i Altaba A. (1999). Sonic hedgehog regulates the growth and patterning of the cerebellum. Development 126, 3089–3100.

20. Damerla, R. R., Cui, C., Gabriel, G. C., Liu, X., Craige, B., Gibbs, B. C., Francis, R., Li, Y., Chatterjee, B., San Agustin, J. T., et al. (2015). Novel Jbts17 mutant mouse model of Joubert syndrome with cilia transition zone defects and cerebellar and other ciliopathy related anomalies. Hum Mol Genet 24, 3994–4005.

21. Dekens, M. P. S., Fontinha, B. M., Gallach, M., Pflügler, S. and Tessmar-Raible, K. (2022). Melanopsin elevates locomotor activity during the wake state of the diurnal zebrafish. EMBO reports 23, e51528.

22. Del Bigio, M. R. (2010). Ependymal cells: biology and pathology. Acta Neuropathol 119, 55–73.

23. DeMars, K. M., Ross, M. R., Starr, A. and McIntyre, J. C. (2023). Neuronal primary cilia integrate peripheral signals with metabolic drives. Frontiers in Physiology 14.

24. Diaz Verdugo, C., Myren-Svelstad, S., Aydin, E., Van Hoeymissen, E., Deneubourg, C., Vanderhaeghe, S., Vancraeynest, J., Pelgrims, R., Cosacak, M. I., Muto, A., et al. (2019). Glia-neuron interactions underlie state transitions to generalized seizures. Nat Commun 10, 3830.

25. Doerre, G. and Malicki, J. (2002). Genetic analysis of photoreceptor cell development in the zebrafish retina. Mech Dev 110, 125–138.

26. Domire, J. S., Green, J. A., Lee, K. G., Johnson, A. D., Askwith, C. C. and Mykytyn, K. (2011). Dopamine receptor 1 localizes to neuronal cilia in a dynamic process that requires the Bardet-Biedl syndrome proteins. Cellular and Molecular Life Sciences 68, 2951–2960.

27. Dreosti, E., Vendrell Llopis, N., Carl, M., Yaksi, E. and Wilson Stephen W. (2014). Left-Right Asymmetry Is Required for the Habenulae to Respond to Both Visual and Olfactory Stimuli. Current Biology 24, 440–445.

28. Duldulao, N. A., Lee, S. and Sun, Z. (2009). Cilia localization is essential for in vivo functions of the Joubert syndrome protein Arl13b/Scorpion. Development 136, 4033–4042.

29. Falk, N., Losl, M., Schroder, N. and Giessl, A. (2015). Specialized Cilia in Mammalian Sensory Systems. Cells 4, 500–519.

30. Faubel, R., Westendorf, C., Bodenschatz, E. and Eichele, G. (2016). Cilia-based flow network in the brain ventricles. Science 353, 176–178.

31. Ferland, R. J., Eyaid, W., Collura, R. V., Tully, L. D., Hill, R. S., Al-Nouri, D., Al-Rumayyan, A., Topcu, M., Gascon, G., Bodell, A., et al. (2004). Abnormal cerebellar development and axonal decussation due to mutations in AHI1 in Joubert syndrome. Nat Genet 36, 1008–1013.

32. Fernandes António M., Fero, K., Arrenberg Aristides B., Bergeron Sadie A., Driever, W. and Burgess Harold A. (2012). Deep Brain Photoreceptors Control Light-Seeking Behavior in Zebrafish Larvae. Current Biology 22, 2042–2047.

33. Fliegauf, M., Benzing, T. and Omran, H. (2007). When cilia go bad: cilia defects and ciliopathies. Nature Reviews Molecular Cell Biology 8, 880–893.

34. Fontinha, B. M., Zekoll, T., Al-Rawi, M., Gallach, M., Reithofer, F., Barker, A. J., Hofbauer, M., Fischer, R. M., von Haeseler, A., Baier, H., et al. (2021). TMT-Opsins differentially modulate medaka brain function in a context-dependent manner. PLoS Biol 19, e3001012.

35. Fore, S., Acuña-Hinrichsen, F., Mutlu, K. A., Bartoszek, E. M., Serneels, B., Faturos, N. G., Chau, K. T. P., Cosacak, M. I., Verdugo, C. D., Palumbo, F., et al. (2020). Functional properties of habenular neurons are determined by developmental stage and sequential neurogenesis. Sci Adv 6.

36. Forsyth, R. and Gunay-Aygun, M. (1993). Bardet-Biedl Syndrome Overview. In GeneReviews(®) (ed. M. P. Adam, G. M. Mirzaa, R. A. Pagon, S. E. Wallace, L. J. H. Bean, K. W. Gripp & A. Amemiya). Seattle (WA): University of Washington, Seattle

37. Copyright © 1993-2023, University of Washington, Seattle. GeneReviews is a registered trademark of the University of Washington, Seattle. All rights reserved.

38. Forsythe, E. and Beales, P. L. (2013). Bardet–Biedl syndrome. European Journal of Human Genetics 21, 8–13.

39. Gabriel, E., Wason, A., Ramani, A., Gooi, L. M., Keller, P., Pozniakovsky, A., Poser, I., Noack, F., Telugu, N. S., Calegari, F., et al. (2016). CPAP promotes timely cilium disassembly to maintain neural progenitor pool. The EMBO Journal 35, 803–819.

40. Guemez-Gamboa, A., Coufal, N. G. and Gleeson, J. G. (2014). Primary cilia in the developing and mature brain. Neuron 82, 511–521.

41. Guo, J., Higginbotham, H., Li, J., Nichols, J., Hirt, J., Ghukasyan, V. and Anton, E. S. (2015). Developmental disruptions underlying brain abnormalities in ciliopathies. Nat Commun 6, 7857.

42. Guo, J., Otis, J. M., Higginbotham, H., Monckton, C., Cheng, J., Asokan, A., Mykytyn, K., Caspary, T., Stuber, G. D. and Anton, E. S. (2017). Primary Cilia Signaling Shapes the Development of Interneuronal Connectivity. Dev Cell 42, 286–300 e284.

43. Guo, J., Otis, J. M., Suciu, S. K., Catalano, C., Xing, L., Constable, S., Wachten, D., Gupton, S., Lee, J., Lee, A., et al. (2019). Primary Cilia Signaling Promotes Axonal Tract Development and Is Disrupted in Joubert Syndrome-Related Disorders Models. Dev Cell 51, 759–774.e755.

44. Hamon, M., Doucet, E., Lefèvre, K., Miquel, M.-C., Lanfumey, L., Insausti, R., Frechilla, D., Del Rio, J. and Vergé, D. (1999). Antibodies and Antisense Oligonucleotide for Probing the Distribution and Putative Functions of Central 5-HT6 Receptors. Neuropsychopharmacology 21, 68S–76S.

45. Hansen, J. N., Rassmann, S., Stüven, B., Jurisch-Yaksi, N. and Wachten, D. (2021). CiliaQ: a simple, open-source software for automated quantification of ciliary morphology and fluorescence in 2D, 3D, and 4D images. Eur Phys J E Soft Matter 44, 18.

46. Higginbotham, H., Eom, T. Y., Mariani, L. E., Bachleda, A., Hirt, J., Gukassyan, V., Cusack, C. L., Lai, C., Caspary, T. and Anton, E. S. (2012). Arl13b in primary cilia regulates the migration and placement of interneurons in the developing cerebral cortex. Dev Cell 23, 925–938.

47. Higginbotham, H., Guo, J., Yokota, Y., Umberger, N. L., Su, C. Y., Li, J., Verma, N., Hirt, J., Ghukasyan, V., Caspary, T., et al. (2013). Arl13b-regulated cilia activities are essential for polarized radial glial scaffold formation. Nat Neurosci 16, 1000–1007.

48. Hilgendorf, K. I., Johnson, C. T. and Jackson, P. K. (2016). The primary cilium as a cellular receiver: organizing ciliary GPCR signaling. Current Opinion in Cell Biology 39, 84–92.

49. Hotz, A. L., Jamali, A., Rieser, N. N., Niklaus, S., Aydin, E., Myren-Svelstad, S., Lalla, L., Jurisch-Yaksi, N., Yaksi, E. and Neuhauss, S. C. F. (2022). Loss of glutamate transporter eaat2a leads to aberrant neuronal excitability, recurrent epileptic seizures, and basal hypoactivity. Glia 70, 196–214.

50. Huangfu, D., Liu, A., Rakeman, A. S., Murcia, N. S., Niswander, L. and Anderson, K. V. (2003). Hedgehog signalling in the mouse requires intraflagellar transport proteins. Nature 426, 83–87.

51. Händel, M., Schulz, S., Stanarius, A., Schreff, M., Erdtmann-Vourliotis, M., Schmidt, H., Wolf, G. and Höllt, V. (1999). Selective targeting of somatostatin receptor 3 to neuronal cilia. Neuroscience 89, 909–926.

52. Insinna, C. and Besharse, J. C. (2008). Intraflagellar transport and the sensory outer segment of vertebrate photoreceptors. Dev Dyn 237, 1982–1992.

53. Jetti, S. K., Vendrell-Llopis, N. and Yaksi, E. (2014). Spontaneous activity governs olfactory representations in spatially organized habenular microcircuits. Curr Biol 24, 434–439.

54. Joubert, M., Eisenring, J. J., Robb, J. P. and Andermann, F. (1969). Familial agenesis of the cerebellar vermis. A syndrome of episodic hyperpnea, abnormal eye movements, ataxia, and retardation. Neurology 19, 813–825.

55. Jurisch-Yaksi, N., Yaksi, E. and Kizil, C. (2020). Radial glia in the zebrafish brain: Functional, structural, and physiological comparison with the mammalian glia. Glia.

56. Kirkby, Lowry A., Sack Georgeann S., Firl, A. and Feller Marla B. (2013). A Role for Correlated Spontaneous Activity in the Assembly of Neural Circuits. Neuron 80, 1129–1144.

57. Kumamoto, N., Gu, Y., Wang, J., Janoschka, S., Takemaru, K., Levine, J. and Ge, S. (2012). A role for primary cilia in glutamatergic synaptic integration of adult-born neurons. Nat Neurosci 15, 399–405, S391.

58. Laurà, R., Magnoli, D., Zichichi, R., Guerrera, M. C., De Carlos, F., Suárez, A. Á., Abbate, F., Ciriaco, E., Vega, J. A. and Germanà, A. (2012). The photoreceptive cells of the pineal gland in adult zebrafish (Danio rerio). Microscopy Research and Technique 75, 359–366.

59. Lee, E. Y., Ji, H., Ouyang, Z., Zhou, B., Ma, W., Vokes, S. A., McMahon, A. P., Wong, W. H. and Scott, M. P. (2010). Hedgehog pathway-regulated gene networks in cerebellum development and tumorigenesis. Proceedings of the National Academy of Sciences 107, 9736–9741.

60. Lepanto, P., Davison, C., Casanova, G., Badano, J. L. and Zolessi, F. R. (2016). Characterization of primary cilia during the differentiation of retinal ganglion cells in the zebrafish. Neural Development 11, 10.

61. Lindsey, B. W., Hall, Z. J., Heuzé, A., Joly, J.-S., Tropepe, V. and Kaslin, J. (2018). The role of neuro-epithelial-like and radial-glial stem and progenitor cells in development, plasticity, and repair. Progress in Neurobiology 170, 99–114.

62. Liu, S., Trupiano, M. X., Simon, J., Guo, J. and Anton, E. S. (2021). The essential role of primary cilia in cerebral cortical development and disorders. Curr Top Dev Biol 142, 99–146.

63. Liu, Y., Nour, M. M., Schuck, N. W., Behrens, T. E. J. and Dolan, R. J. (2022). Decoding cognition from spontaneous neural activity. Nature Reviews Neuroscience 23, 204–214.

64. Louvi, A. and Grove Elizabeth A. (2011). Cilia in the CNS: The Quiet Organelle Claims Center Stage. Neuron 69, 1046–1060.

65. Masek, M., Zang, J., Mateos, J. M., Garbelli, M., Ziegler, U., Neuhauss, S. C. F. and Bachmann-Gagescu, R. (2023). Studying the morphology, composition and function of the photoreceptor primary cilium in zebrafish. Methods Cell Biol 175, 97–128.

66. McClintock, T. S., Khan, N., Xie, C. and Martens, J. R. (2020). Maturation of the Olfactory Sensory Neuron and Its Cilia. Chem Senses 45, 805–822.

67. Menco, B. P. M., Cunningham, A. M., Qasba, P., Levy, N. and Reed, R. R. (1997). Putative odour receptors localize in cilia of olfactory receptor cells in rat and mouse: a freeze-substitution ultrastructural study. Journal of Neurocytology 26, 691–706.

68. Mitchison, H. M. and Valente, E. M. (2017). Motile and non-motile cilia in human pathology: from function to phenotypes. J Pathol 241, 294–309.

69. Mueller, T. (2012). What is the Thalamus in Zebrafish? Frontiers in Neuroscience 6.

70. Myren-Svelstad, S., Jamali, A., Ophus, S. S., D’gama, P. P., Ostenrath, A. M., Mutlu, A. K., Hoffshagen, H. H., Hotz, A. L., Neuhauss, S. C. F., Jurisch-Yaksi, N., et al. (2022). Elevated photic response is followed by a rapid decay and depressed state in ictogenic networks. Epilepsia 63, 2543–2560.

71. Nachury, M. V. (2014). How do cilia organize signalling cascades? Philos Trans R Soc Lond B Biol Sci 369.

72. Niklaus, S., Cadetti, L., Vom Berg-Maurer, C. M., Lehnherr, A., Hotz, A. L., Forster, I. C., Gesemann, M. and Neuhauss, S. C. F. (2017). Shaping of Signal Transmission at the Photoreceptor Synapse by EAAT2 Glutamate Transporters. eNeuro 4.

73. Ohki, K., Chung, S., Ch’ng, Y. H., Kara, P. and Reid, R. C. (2005). Functional imaging with cellular resolution reveals precise micro-architecture in visual cortex. Nature 433, 597–603.

74. Olstad, E. W., Ringers, C., Hansen, J. N., Wens, A., Brandt, C., Wachten, D., Yaksi, E. and Jurisch-Yaksi, N. (2019). Ciliary Beating Compartmentalizes Cerebrospinal Fluid Flow in the Brain and Regulates Ventricular Development. Curr Biol 29, 229–241.e226.

75. Omori, Y., Zhao, C., Saras, A., Mukhopadhyay, S., Kim, W., Furukawa, T., Sengupta, P., Veraksa, A. and Malicki, J. (2008). Elipsa is an early determinant of ciliogenesis that links the IFT particle to membrane-associated small GTPase Rab8. Nat Cell Biol 10, 437–444.

76. Paridaen, J. T. and Huttner, W. B. (2014). Neurogenesis during development of the vertebrate central nervous system. EMBO Rep 15, 351–364.

77. Parisi, M. and Glass, I. (1993). Joubert Syndrome. In GeneReviews(®) (ed. M. P. Adam, G. M. Mirzaa, R. A. Pagon, S. E. Wallace, L. J. H. Bean, K. W. Gripp & A. Amemiya). Seattle (WA): University of Washington, Seattle Copyright © 1993-2023, University of Washington, Seattle. GeneReviews is a registered trademark of the University of Washington, Seattle. All rights reserved.

78. Park, S. M., Jang, H. J. and Lee, J. H. (2019). Roles of Primary Cilia in the Developing Brain. Front Cell Neurosci 13, 218.

79. Plotnikova, O. V., Pugacheva, E. N. and Golemis, E. A. (2009). Primary cilia and the cell cycle. Methods Cell Biol 94, 137–160.

80. Preibisch, S., Saalfeld, S. and Tomancak, P. (2009). Globally optimal stitching of tiled 3D microscopic image acquisitions. Bioinformatics 25, 1463–1465.

81. Prigent, C. and Dimitrov, S. (2003). Phosphorylation of serine 10 in histone H3, what for? Journal of Cell Science 116, 3677–3685.

82. Rachel, R. A., Yamamoto, E. A., Dewanjee, M. K., May-Simera, H. L., Sergeev, Y. V., Hackett, A. N., Pohida, K., Munasinghe, J., Gotoh, N., Wickstead, B., et al. (2015). CEP290 alleles in mice disrupt tissue-specific cilia biogenesis and recapitulate features of syndromic ciliopathies. Hum Mol Genet 24, 3775–3791.

83. Recher, G., Jouralet, J., Brombin, A., Heuzé, A., Mugniery, E., Hermel, J.-M., Desnoulez, S., Savy, T., Herbomel, P., Bourrat, F., et al. (2013). Zebrafish midbrain slow-amplifying progenitors exhibit high levels of transcripts for nucleotide and ribosome biogenesis. Development 140, 4860–4869.

84. Reiten, I., Uslu, F. E., Fore, S., Pelgrims, R., Ringers, C., Diaz Verdugo, C., Hoffman, M., Lal, P., Kawakami, K., Pekkan, K., et al. (2017). Motile-Cilia-Mediated Flow Improves Sensitivity and Temporal Resolution of Olfactory Computations. Curr Biol 27, 166–174.

85. Reiter, J. F. and Leroux, M. R. (2017). Genes and molecular pathways underpinning ciliopathies. Nature reviews. Molecular cell biology 18, 533–547.

86. Ringers, C., Olstad, E. W. and Jurisch-Yaksi, N. (2020). The role of motile cilia in the development and physiology of the nervous system. Philosophical Transactions of the Royal Society B: Biological Sciences 375, 20190156.

87. Robles, E., Laurell, E. and Baier, H. (2014). The Retinal Projectome Reveals Brain-Area-Specific Visual Representations Generated by Ganglion Cell Diversity. Current Biology 24, 2085–2096.

88. Romano, S. A., Pérez-Schuster, V., Jouary, A., Boulanger-Weill, J., Candeo, A., Pietri, T. and Sumbre, G. (2017). An integrated calcium imaging processing toolbox for the analysis of neuronal population dynamics. PLoS Comput Biol 13, e1005526.

89. Rosso, S., Inestrosa, N. and Rosso, S. (2013). WNT signaling in neuronal maturation and synaptogenesis. Frontiers in Cellular Neuroscience 7.

90. Rusterholz, T. D. S., Hofmann, C. and Bachmann-Gagescu, R. (2022). Insights Gained From Zebrafish Models for the Ciliopathy Joubert Syndrome. Front Genet 13, 939527.

91. Samarut, É., Swaminathan, A., Riché, R., Liao, M., Hassan-Abdi, R., Renault, S., Allard, M., Dufour, L., Cossette, P., Soussi-Yanicostas, N., et al. (2018). γ-Aminobutyric acid receptor alpha 1 subunit loss of function causes genetic generalized epilepsy by impairing inhibitory network neurodevelopment. Epilepsia 59, 2061–2074.

92. Sawamoto, K., Wichterle, H., Gonzalez-Perez, O., Cholfin, J. A., Yamada, M., Spassky, N., Murcia, N. S., Garcia-Verdugo, J. M., Marin, O., Rubenstein, J. L., et al. (2006). New neurons follow the flow of cerebrospinal fluid in the adult brain. Science 311, 629–632.

93. Sengupta, P., Chou, J. H. and Bargmann, C. I. (1996). odr-10 Encodes a Seven Transmembrane Domain Olfactory Receptor Required for Responses to the Odorant Diacetyl. Cell 84, 899–909.

94. Sheu, S.-H., Upadhyayula, S., Dupuy, V., Pang, S., Deng, F., Wan, J., Walpita, D., Pasolli, H. A., Houser, J., Sanchez-Martinez, S., et al. (2022). A serotonergic axon-cilium synapse drives nuclear signaling to alter chromatin accessibility. Cell 185, 3390–3407.e3318.

95. Stoufflet, J. and Caillé, I. (2022). The Primary Cilium and Neuronal Migration. Cells 11.

96. Stoufflet, J., Chaulet, M., Doulazmi, M., Fouquet, C., Dubacq, C., Métin, C., Schneider-Maunoury, S., Trembleau, A., Vincent, P. and Caillé, I. (2020). Primary cilium-dependent cAMP/PKA signaling at the centrosome regulates neuronal migration. Sci Adv 6.

97. Suciu, S. K. and Caspary, T. (2021). Cilia, neural development and disease. Seminars in Cell & Developmental Biology 110, 34–42.

98. Tu, H.-Q., Li, S., Xu, Y.-L., Zhang, Y.-C., Li, P.-Y., Liang, L.-Y., Song, G.-P., Jian, X.-X., Wu, M., Song, Z.-Q., et al. (2023). Rhythmic cilia changes support SCN neuron coherence in circadian clock. Science 380, 972–979.

99. Tucker, R. W., Pardee, A. B. and Fujiwara, K. (1979). Centriole ciliation is related to quiescence and DNA synthesis in 3T3 cells. Cell 17, 527–535.

100. Uddin, L. Q. (2020). Bring the Noise: Reconceptualizing Spontaneous Neural Activity. Trends in Cognitive Sciences 24, 734–746.

101. Vigh, B., Manzano, M. J., Zádori, A., Frank, C. L., Lukáts, A., Röhlich, P., Szél, A. and Dávid, C. (2002). Nonvisual photoreceptors of the deep brain, pineal organs and retina. Histol Histopathol 17, 555–590.

102. Vladimirov, N., Mu, Y., Kawashima, T., Bennett, D. V., Yang, C. T., Looger, L. L., Keller, P. J., Freeman, J. and Ahrens, M. B. (2014). Light-sheet functional imaging in fictively behaving zebrafish. Nat Methods 11, 883–884.

103. Wachten, D. and Mick, D. U. (2021). Signal transduction in primary cilia – analyzing and manipulating GPCR and second messenger signaling. Pharmacology & Therapeutics 224, 107836.

104. Wallace, V. A. (1999). Purkinje-cell-derived Sonic hedgehog regulates granule neuron precursor cell proliferation in the developing mouse cerebellum. Curr Biol 9, 445–448.

105. Wang, W., Shiraishi, R. and Kawauchi, D. (2022). Sonic Hedgehog Signaling in Cerebellar Development and Cancer. Front Cell Dev Biol 10, 864035.

106. Wang, Y., Bernard, A., Comblain, F., Yue, X., Paillart, C., Zhang, S., Reiter, J. F. and Vaisse, C. (2021). Melanocortin 4 receptor signals at the neuronal primary cilium to control food intake and body weight. The Journal of Clinical Investigation 131.

107. Wechsler-Reya, R. J. and Scott, M. P. (1999). Control of neuronal precursor proliferation in the cerebellum by Sonic Hedgehog. Neuron 22, 103–114.

108. Yaksi, E., Jamali, A., Diaz Verdugo, C. and Jurisch-Yaksi, N. (2021). Past, present and future of zebrafish in epilepsy research. Febs j 288, 7243–7255.

109. Youn, Y. H. and Han, Y. G. (2018). Primary Cilia in Brain Development and Diseases. Am J Pathol 188, 11–22.

110. Zhang, B.-b., Yao, Y.-y., Zhang, H.-f., Kawakami, K. and Du, J.-l. (2017). Left Habenula Mediates Light-Preference Behavior in Zebrafish via an Asymmetrical Visual Pathway. Neuron 93, 914–928.e914.

111. Zhu, J., Wang, H.-T., Chen, Y.-R., Yan, L.-Y., Han, Y.-Y., Liu, L.-Y., Cao, Y., Liu, Z.-Z. and Xu, H. A. (2020). The Joubert Syndrome Gene arl13b is Critical for Early Cerebellar Development in Zebrafish. Neuroscience Bulletin 36, 1023–1034.

