## Supplemental Figures for "Loss of cilia after neurulation impacts brain development and neuronal activity in larval zebrafish"

**Supplementary figure 1:**

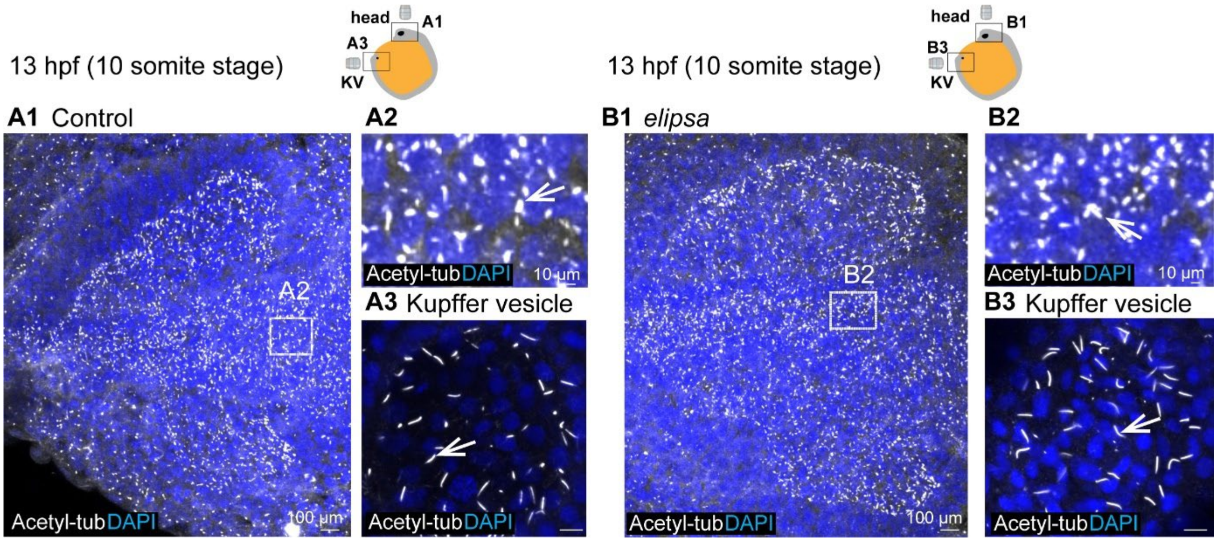


**Cilia are present at 10 somites stage in *elipsa* mutants.**

**(A1-A2** and **B1-B2)** Acetylated tubulin staining of 13 hpf zebrafish larvae showing presence of cilia (white) in the head region in both control **(A1)** n=12 and *elipsa* mutant **(B1)** n=5**. (A2, B2)** Insets drawn in the region near the neural tube to indicate presence of cilia in control **(A2)** and mutant **(B2)**. **(A3, B3)** Kupffer's vesicle labeling with acetylated tubulin (white) show similar numbers of cilia at 13hpf in control **(A3)** n=12 and mutant **(B3)** n=5**.**

**Supplementary figure 2:**

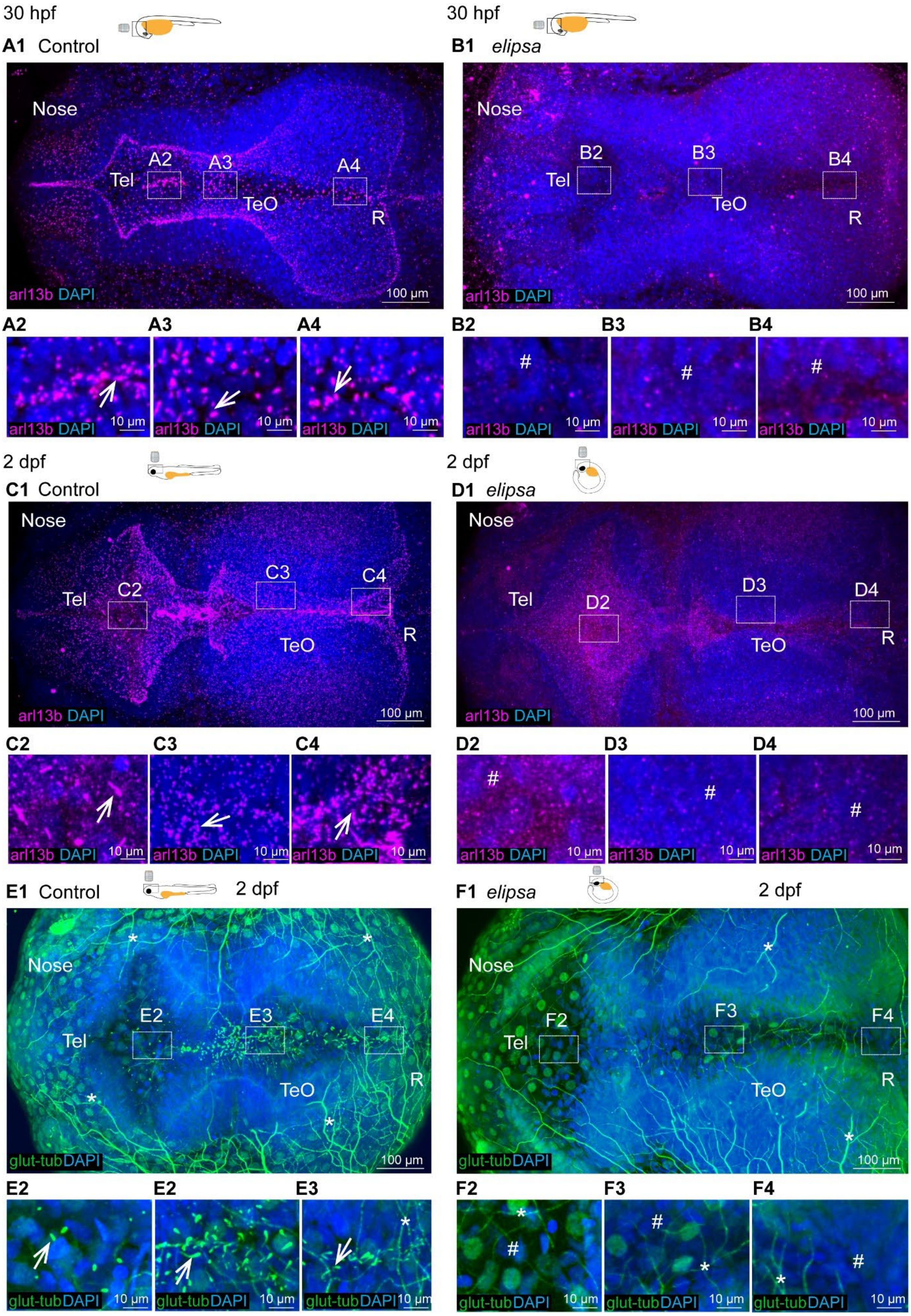

**Cilia are lost in the brain of 30hpf-2dpf *elipsa* mutant embryos.**

**(A1-A4** and **B1-B4)** Staining of 30hpf zebrafish larvae with arl3b antibody to stain all cilia in the brain, n=9 controls and 4 mutants. **A1** At 30 hpf, arl13b stained cilia are located in various brain regions, represented by insets drawn in different brain regions **(A2-A4). (B1)** Loss of cilia in the entire brain of *elipsa*, represented by insets drawn in different regions **(B2-B4).** (**C1-C4** and **D1-D4**) Staining of 2dpf brains with arl13b antibody to stain cilia in the brain, n=3 controls and 3 mutants. **(D1)** Loss of cilia in *elipsa* mutant brains, as shown by insets drawn in different regions **(D2-D4).** (**E1-E4** and **F1-F4**) Glutamylated tubulin staining for motile cilia at 2dpf, n=3 controls and 4 mutants. **(E1) S**ingle glutamylated tubulin-positive cilia are located in the forebrain on the dorsal roof and ventral part of the tectal/diencephalic ventricle and in the rhombencephalon, as indicated by insets drawn in the telencephalon **(E2)**, optic tectum**(E3)** and rhombencephalon **(E4). (F1)** Loss of *elipsa* leads to loss of glutamylated tubulin-positive cilia in the brain, represented by insets drawn in different regions **(F2-F4).** Tel, Telencephalon; Teo, Optic Tectum; R, Rhombencephalon. Cilia loss is indicated by # and nonspecific signal from glutamylated tubulin antibody by *.

**Supplementary figure 3:**

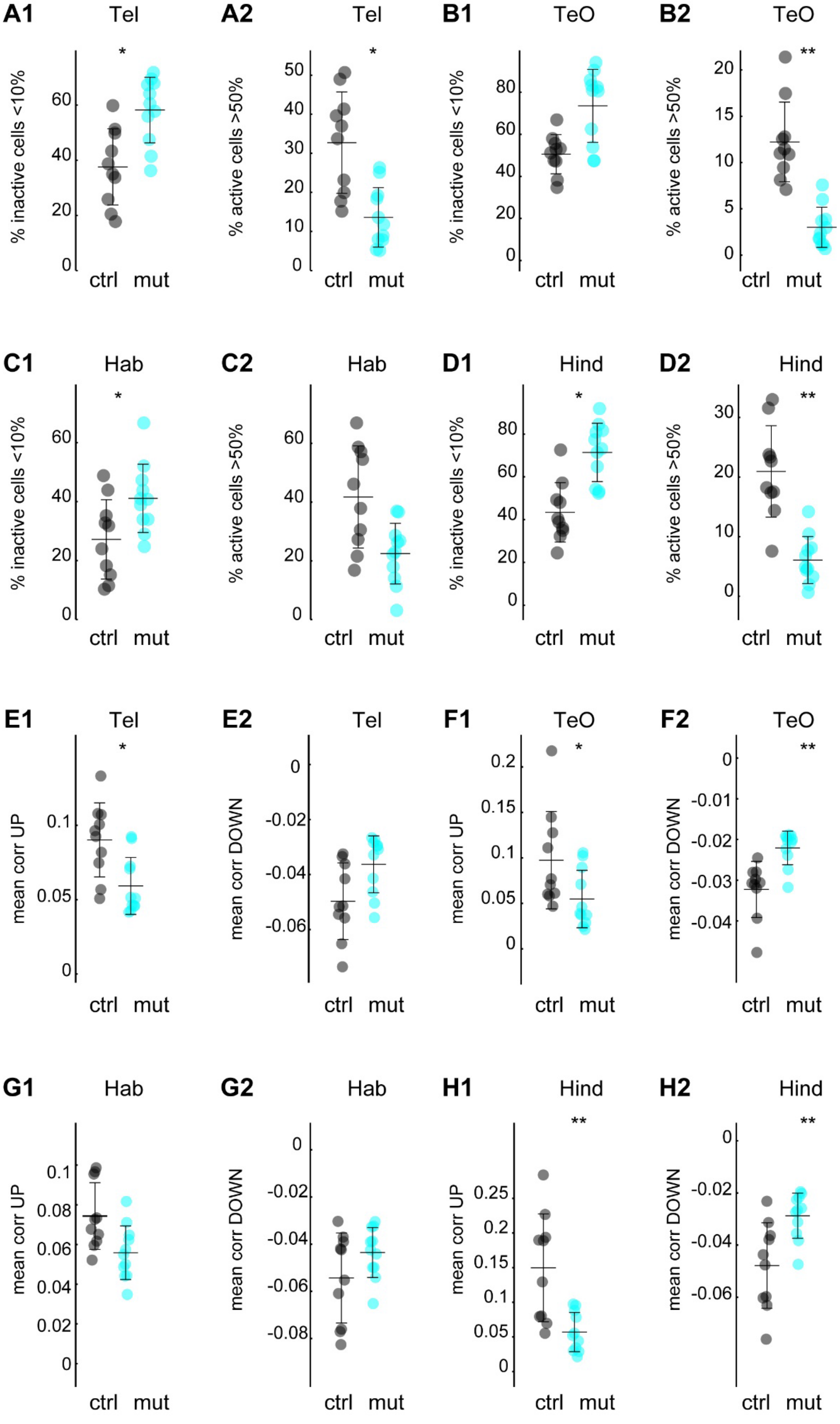

**Reduced spontaneous activity and correlation between neighboring neurons in the brain of cilia mutants.**
**(A1-D2)** Quantification of inactive and active cells for the **(A1-A2)** telencephalon, **(B1-B2)** optic tectum and **(C1-C2)** habenula and **(D1-D2)** hindbrain regions, controls in black and *elipsa* mutant in cyan. N=10 controls and 11 mutants. **(E1-H2)** Quantification of mean positive (UP) and negative (DOWN) correlation for cells located within 60 µm distance in the **(E1-E2)** telencephalon, **(F1-F2)** optic tectum, **(G1-G2)** habenula and **(H1-H2)** hindbrain regions, controls in black and *elipsa* mutant in cyan. N=10 controls and 11 mutants. Statistical significance by Wilcoxon Rank Sum test, * $p<0.05$, ** $p<0.01$. Tel, Telencephalon; Teo, Optic Tectum; Hab, Habenula; Hind, Hindbrain.

**<u>Supplementary table 1:</u>**

RNA sequencing results obtained from 4 dpf control and *elipsa* mutant larvae.
